# GSNOR-dependent nitric oxide homeostasis shapes methylome persistence after multi-generational exposure to future climate conditions in *Arabidopsis thaliana*

**DOI:** 10.64898/2026.11.09.765246

**Authors:** R Behl, B Thanh Vo, A Ghirardo, A Albert, E Mattes, L Gößl, JP Schnitzler, F Johannes, C Lindermayr

## Abstract

Environmental stress can modify plant DNA methylation, but the roles of nitric oxide homeostasis in methylome recovery and multigenerational persistence remain unclear. We used whole-genome bisulfite sequencing to compare *Arabidopsis thaliana* wild type and the GSNOR-deficient mutant (*gsnor1-3*) exposed to drought, elevated CO_2_, ozone, heat, or their combination during generation G1 to G3, followed by two generations under control conditions. Methylomes from G1, G3, and G5 were analysed for global divergence, differentially methylated regions, genomic annotation, and methylation trajectories and were integrated with matched RNA-seq and independent datasets. Control methylomes of *gsnor1-3* were already distinct from those of the wild type, and the mutant accumulated substantially more DMRs, with a strong hypermethylation bias. In *gsnor1-3*, 44.1 to 45.7% of G1 CG DMRs retained their direction through G3 and G5, compared with 0.09 to 1.24% in wild type; persistent CHG and CHH DMRs were also markedly more abundant. Persistent regions occurred at genes, promoters, and transposable elements and were associated with broader expression responses in the mutant. Independent datasets supported a subset of these regions, including directional agreement for 211 of 292 published GSNOR-associated DMRs. These results indicate that GSNOR-dependent nitric oxide homeostasis contributes to methylome stability and resetting after repeated environmental exposure.

**PLAIN LANGUAGE SUMMARY:** Plants can retain molecular traces of environmental stress, but it is unclear why some traces disappear while others persist in later generations. We exposed wild-type Arabidopsis plants and a GSNOR-deficient mutant to drought, elevated carbon dioxide, ozone, heat, or their combination for three generations. GSNOR regulates nitric oxide levels and has important function in stress response. After stress exposure, plants were grown under control conditions for two generations. The mutant has more differentially methylated regions than wild-type plants even under control conditions, developed many more changes under stress conditions, and retained a much larger proportion during recovery. Many retained changes occurred near genes and mobile DNA elements and coincided with altered gene activity. Our results suggest that GSNOR-dependent regulation of nitric oxide helps plants restore their DNA methylation landscape after repeated environmental stress.

Graphical abstract:
*Created in BioRender.com*

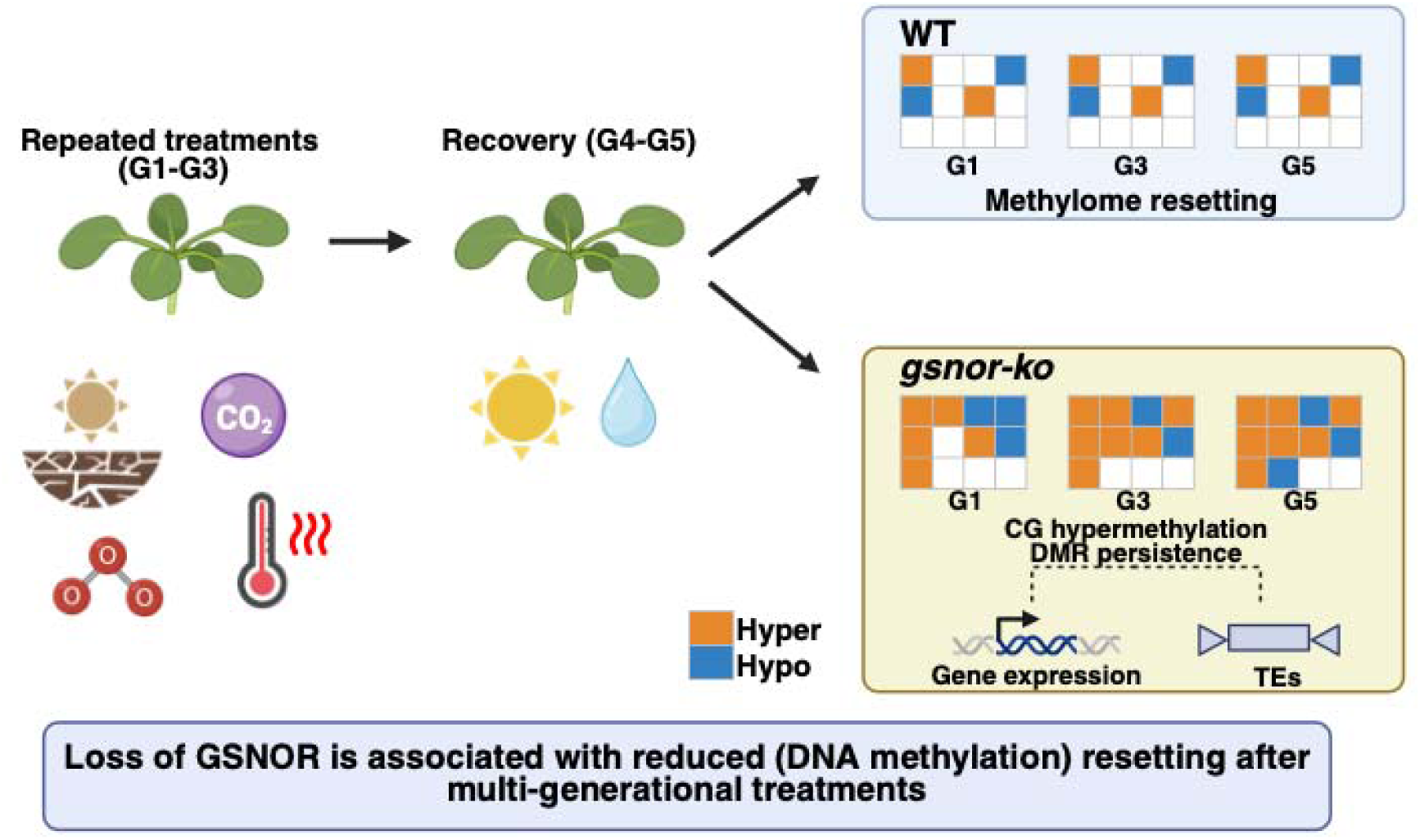

## 1. INTRODUCTION

Plants are constantly subjected to environmental fluctuations that limit their growth, reproduction, and survival (Joseph et al., 2025; Zhang et al., 2023). Due to anthropogenic climate change, the frequency and intensity of high-temperature events have increased, and, in many areas, so have agricultural and ecological droughts. This means that plants are subjected to a range of stresses, including changes in atmospheric CO_2_ concentration, increases in tropospheric ozone, and other environmental disturbances (IPCC, 2023a, 2023b). However, these different factors rarely occur in isolation, and plants may therefore be exposed to more than one stress at a time, one after another or repeatedly. The way in which plants respond to a combination of stresses cannot always be deduced from their responses to each stress on its own. In *Arabidopsis thaliana*, for example, simultaneous drought and heat stress led to a specific physiological and transcriptional response, involving hundreds of transcripts not activated by either stress alone (Behl et al., 2026; Rizhsky et al., 2004). Similar results have been found in studies involving sequential stress, in which a prior stress has left detectable transcriptional marks and altered either the timing or the extent of the response to a subsequent stress (Coolen et al., 2016). It is therefore essential to understand how plants respond to repeated and interacting environmental conditions if we are to predict their performance in future climates and identify the molecular processes that contribute to resilience.

Plants undergo changes in transcription, along with changes in hormone signalling, metabolism, redox regulation, growth, and resource distribution, in response to stress (Fernandes & Ghag, 2022). Even though several of the responses decrease following the removal of the stress, prior exposure can affect the rate, extent or direction of the response to a subsequent stress. This is generally referred to as stress priming or stress memory (Crisp et al., 2016; Sani et al., 2013). For instance, repeated dehydration in Arabidopsis leads to the formation of different categories of transcriptional memory genes, some of which are induced more strongly, others less strongly, and in some cases show altered response patterns upon later exposure (Ding et al., 2012; Ding et al., 2013). In certain cases involving dehydration-memory gene loci, transcription returns to baseline levels during recovery, even though elevated H3K4me3 and phosphorylated RNA polymerase II remain associated with the genes, indicating that a molecular record of the previous stress can persist even after transcription has been restored (Ding et al., 2012). Heat-stress memory is also linked to sustained H3K4 methylation and the activity of specific heat-shock transcription factors, indicating that chromatin states play a role in maintaining the transcriptional competence induced by stress (Liu et al., 2018). However, it is not always beneficial to maintain a stress-related state, since effective recovery also involves switching off the protective programmes and restoring growth. Plant performance thus relies on a balance between molecular memory and the resetting that occurs after stress is removed (Crisp et al., 2016).

Epigenetic mechanisms offer a possible molecular basis for this balance by altering how genes are expressed. Cytosine DNA methylation is particularly significant in plants and occurs in the CG, CHG, and CHH sequence contexts, with H representing A, C, or T (Abdul Aziz & Masmoudi, 2026; Johannes & Schmitz, 2019; Law & Jacobsen, 2010). The pathways responsible for establishing and maintaining these three contexts differ in part (Zhang et al., 2018). CG methylation is mainly kept up by METHYLTRANSFERASE 1, while CHG methylation is mostly maintained by CHROMOMETHYLASE 3 (Cao & Jacobsen, 2002; Kankel et al., 2003). The DOMAINS REARRANGED METHYLTRANSFERASE proteins are involved in de novo methylation and play a central role in RNA-directed DNA methylation, and CHROMOMETHYLASE 2 makes a major contribution to CHH methylation in heterochromatin that is enriched in H3K9me2 (Cao & Jacobsen, 2002; Stroud et al., 2014). Together, these pathways help to limit the activity of transposable elements (TE), preserve genome stability, and create both transcriptionally permissive and repressive chromatin environments (Liang et al., 2022; Stroud et al., 2014). Since CG and CHG methylation occur at symmetric sites in the DNA sequence, it can be copied during DNA replication, whereas asymmetric CHH methylation must be re-established in each cell generation. Stress-induced methylation changes in these contexts may therefore vary in terms of stability, location in the genome, and ability to last during recovery or from one generation to the next (Boyko et al., 2010; Johannes & Schmitz, 2019; Law & Jacobsen, 2010; Sani et al., 2013; Shahryary, Symeonidi, et al., 2020).

Environmental stresses can cause changes in DNA methylation at genes, promoters, TEs, and other regulatory regions, but it is still unclear to what extent these changes reflect stable stress memory. Some experiments have identified methylation states that are responsive to stress and linked to either changes in gene expression or phenotypic changes in the offspring. For instance, in *Arabidopsis thaliana*, repeated exposure to hyperosmotic stress led to methylation changes at epigenetically labile sites and resulted in a short-term, mainly maternally inherited memory that was widely reset following removal of the stress (Wibowo et al., 2016, 2018). In other cases, DNA methylation and small RNA-dependent responses have been observed in the direct offspring of stressed plants (Boyko et al., 2010).

Redox signalling could be a significant yet underexplored factor in these methylome dynamics. Nitric oxide is a redox-active signalling molecule that primarily alters protein function via S-nitrosylation of cysteine residues. S-nitrosoglutathione reductase regulates this signalling pathway by breaking down S-nitrosoglutathione and, as a result, influencing the amount of cellular S-nitrosothiols and protein S-nitrosylation (Behl et al., 2026; Feechan et al., 2005; Rudolf et al., 2021). When the GSNOR1 function is lost in *Arabidopsis,* there is a buildup and improper regulation of S-nitrosothiols, along with effects on plant development, fertility, thermotolerance, hormone signalling, and immunity (Feechan et al., 2005; Lee et al., 2008; Shi et al., 2015). The *gsnor1-3* null mutant is thus a genetically defined system for investigating how disturbed nitric oxide homeostasis affects plant responses to repeated environmental stress.

Some of the findings provide further evidence of a mechanistic link between nitric oxide homeostasis and epigenetic regulation. Nitric oxide can affect histone acetylation by modulating the activity of histone deacetylases, and GSNOR-dependent changes in nitric oxide levels have been linked to genome-wide alterations in histone modifications and stress-related transcription (Ageeva-Kieferle et al., 2021; Behl et al., 2026). The chromatin responses that are dependent on HDA6 are affected by both GSNOR and nitric oxide, while the S-nitrosylation of HDA19 controls its association with chromatin, its histone deacetylation activity, and its role in tolerating oxidative stress. These findings show that protein S-nitrosylation acts as a redox-sensing mechanism which can alter chromatin activity (Ageeva-Kieferle et al., 2021; Lindermayr et al., 2020; Rudolf et al., 2021; Zheng et al., 2023). It remains largely unknown whether a disruption in nitric oxide homeostasis also affects the establishment, maintenance, or resetting of stress-associated DNA methylation patterns over several generations.

In this study, we performed Whole-Genome Bisulfite Sequencing (WGBS) of a multigenerational experiment to examine how repeated exposure to climate change-related stress and subsequent recovery periods affected the DNA methylomes of both wild-type and *gsnor-ko* individuals. Methylation responses in the CG, CHG, and CHH contexts were analysed across generations, and differentially methylated regions (DMRs) linked to single and combined environmental treatments were identified. We then separated the areas that exhibited persistent hypermethylation or hypomethylation from those that reversed their methylation direction or were no longer detectable after removal of the stress, and examined how these regions were distributed among genes, promoters, TEs, and intergenic regions. We proposed that the majority of methylation changes associated with stress would be reversed after two generations under control conditions, while a smaller group would either persist or switch in direction. We also suggested that disrupting GSNOR-dependent nitric oxide homeostasis would modify both the degree and the genomic location of methylation persistence and resetting. By investigating these changes genome-wide, the present study addresses whether redox homeostasis contributes to the stability of stress-induced epigenomic states across successive generations of exposure and recovery.

## 2. MATERIALS AND METHODS

### 2.1 Plant material and multigenerational climate-stress experiment

The multigenerational climate-stress experiment was conducted as described in (Behl et al., 2026). Wild-type (WT) *Arabidopsis thaliana* Col-0 and *gsnor-ko* plants were grown under control, drought, elevated CO_2_, elevated ozone (O_3_), warm climate (heat), and combined treatments across five successive generations. Treatments were applied during the first three generations (G1-G3). Later generations (G4-G5) were maintained under ambient control conditions to assess recovery and transgenerational effects. Rosette leaves were collected from 4.5-week-old plants between 09:00 and 11:00. For this study, leaf material from G1, G3, and G5 was used for WGBS to investigate stress-induced, inherited, reset, and genotype-dependent DNA methylation states (Supplemental Figure S1 and Supplemental File S1).

### 2.2 Whole-Genome Bisulfite Sequencing

Genomic DNA was extracted from 65–70 mg of ground leaf tissue using a CTAB-based protocol (Healey et al., 2014). DNA quantity and quality were assessed using NanoDrop spectrophotometry and Qubit fluorometry. DNA was sheared to an average fragment size of about 350 bp using the Covaris R230 instrument. Libraries were prepared with NEBNext Ultra II DNA Library Prep Kit (E7645S, NEB, USA). Bisulfite conversion was performed using the EZ-96 DNA Methylation-Gold MagPrep Kit (D5042, Zymo Research, USA). Fragmented DNA was end-repaired, A-tailed, ligated to methylated adapters, size-selected, bisulfite converted, and PCR-amplified using KAPA HiFi HS Uracil+ polymerase (Roche; Cat. No. 07959050001).

WGBS libraries were sequenced by Novogene (UK) Co., Ltd. on the NovaSeq X Plus platform using 150 bp paired-end sequencing. Approximately 7 Gb of sequencing data were generated per sample. Three biological replicates were used for the control samples, and two for each treatment in G1, G3, and G5.

### 2.3 WGBS read processing and methylation calling

WGBS reads were processed using the MethylStar pipeline, which integrates adapter and quality trimming with Trimmomatic, bisulfite-aware alignment with Bismark, and cytosine-level methylation calling with Methimpute, as described previously (Shahryary, Hazarika, et al., 2020; Taudt et al., 2018). Reads were aligned to the *Arabidopsis thaliana* TAIR10 reference genome. DNA methylation was analysed separately in CG, CHG, and CHH sequence contexts.

Differentially methylated regions (DMRs) were identified using jDMRgrid (Behl et al., 2025; Hazarika et al., 2022). The genome was partitioned into 200-bp sliding windows with a 50-bp step size. Only windows containing at least 10 cytosines in the respective methylation context were retained. Methylation region calls were generated for each sample and used to construct region-call matrices for pairwise comparisons between control and stress conditions. Non-epipolymorphic patterns were removed, replicate consensus filtering was applied, and adjacent significant windows were merged to obtain final DMRs.

DMRs representing treatment responses were identified by comparing each treatment with the WT control of the respective generation. Direct genotype contrasts were additionally performed by comparing *gsnor-ko* and WT under the same treatment and generation. DMRs were classified as hypermethylated or hypomethylated relative to the WT control.

### 2.4 Global methylation divergence and AlphaBeta analysis

Genome-wide methylation was analysed separately for each sequence context using biological-replicate methylation estimates. Within each generation, a two-factor ANOVA was fitted with genotype, treatment and their interaction as fixed effects. Post-hoc comparisons were performed using the Dunnett test, with p-values adjusted using the Benjamini–Hochberg procedure.

To evaluate whether methylation changes accumulated across generations and to quantify methylation stability, spontaneous CG epimutation rates were estimated for WT and *gsnor-ko* using the AlphaBeta R package (v1.10.0) (Shahryary, Symeonidi, et al., 2020). Following published approaches (Hazarika et al., 2022; van der Graaf et al., 2015), CG methylation gain (α) and loss (β) rates were inferred from multigenerational WGBS data using known pedigree structures. Confidence intervals were computed as implemented in AlphaBeta.

### 2.5 Annotation of DMRs

To assess the fate of stress-associated methylation changes across generations, overlapping DMRs from G1, G3, and G5 were merged into consensus genomic regions. For each consensus region, methylation state was assigned in each generation as hypermethylated, hypomethylated, mixed/both, or absent. Based on the G1–G3–G5 methylation-state pattern, each region was classified into trajectory classes, including acute/generation-specific, stress-maintained-then-reset, persistent, recurrent, G3/G5-emergent, and switcher DMRs. These trajectory classes were used to distinguish methylation changes that were reset after stress withdrawal from those that persisted, reappeared, or changed direction across generations.

Gene, TE, and genome annotation files were obtained from TAIR10. Promoters were defined as 1 kb upstream of annotated gene transcription start sites. DMRs were annotated according to their overlap with gene bodies, promoters, TEs, and intergenic regions. DMR-associated genes and TEs were extracted for downstream analyses of overlap, trajectories, and enrichment. Where DMRs overlapped more than one annotation class, multiple annotation labels were retained for downstream feature-specific summaries.

### 2.6 TEs and genes in their proximity

To determine whether persistent methylation marks were preferentially associated with specific TE classes, TE-overlapping DMRs were mapped to TAIR10 TE family and superfamily annotations. TE superfamilies included, where present, Helitron, Gypsy, Copia, MuDR, HAT, En-Spm, Mariner/Tc1, LINE, and SINE classes. Persistent DMR-associated TEs were compared against all DMR-associated TEs and, separately, against all annotated TAIR10 TEs.

To assess potential regulatory relationships between TE-associated methylation changes and nearby gene expression, TE-DMRs were paired with genes located within approximately ±1.2 kb (Behl et al., 2025). This distance threshold was selected to focus on the closest quartile of TE–gene pairs. For each TE–gene pair, the TE-DMR methylation state and the corresponding gene expression state were combined to define classes: TE hypermethylation with gene downregulation; TE hypermethylation with gene upregulation; TE hypomethylation with gene upregulation; TE hypomethylation with gene downregulation; and TE hyper/hypomethylation with unchanged gene expression. Persistent TE-DMRs and switcher TE-DMRs were also compared with gene-expression trajectories across G1, G3, and G5. These analyses were used to identify candidate TE-linked regulatory relationships and were interpreted as associations rather than direct evidence of causality.

### 2.7 Candidate regulatory gene and Integration with WGBS and RNA-seq data from the previous manuscripts

RNA-seq results from the same experimental system were obtained from our previous study (Behl et al., 2026). Because the RNA-seq and WGBS datasets were generated from the same genotypes, stress treatments, and generations, RNA-seq results were integrated with DMRs and gene annotations to assess methylation–expression relationships. Additionally, previously published environmental stress-associated DMRs were compared with DMRs identified in this study to establish a link between stress-induced DMRs and their inheritance (Behl et al., 2025; Rudolf et al., 2021).

### 2.8 Statistical analysis and visualization

All statistical analyses were performed in R/RStudio. Data processing and visualisation were performed using R packages including data.table, dplyr, tidyr, ggplot2, GenomicRanges, IRanges, rtracklayer, and ggalluvial. Final figure assembly and graphical adjustments were performed in Inkscape.

### 2.9 AI usage in Data analysis

For data analysis, an AI tool (ChatGPT 5.6) was used for programming in R (2024.12.1+563). After using this tool/service, the author(s) reviewed and edited the codes as needed and take(s) full responsibility for the content of the published article.

## 3. RESULTS

We investigated genome-wide DNA methylation dynamics in WT and *gsnor-ko* during repeated environmental exposure in G1–G3 followed by two generations of recovery in G4 and G5. The experimental design and growth conditions are described in Methods and Supplemental Figure S1.

### 3.1 Loss of GSNOR increases DMR abundance and promotes hypermethylation across generations

We examined the methylomes of control samples to determine whether genotype-dependent differences were present in the absence of environmental treatment. Principal Component Analysis (PCA) separated WT and *gsnor-ko* control samples in all three methylation contexts (Supplemental Figure S2). The biological replicates were generally grouped by genotype, especially in the CG context, indicating consistent methylation profiles. The separation between WT and *gsnor-ko* controls demonstrated that loss of GSNOR was associated with an altered baseline methylome prior to treatment-dependent changes.

Genome-wide average methylation differed among methylation contexts, generations, genotypes, and treatments (Supplemental Figure S3). Absolute changes in average methylation were relatively modest compared with the regional differences identified by DMR analysis. Nevertheless, *gsnor-ko* generally displayed higher methylation than WT across several treatment conditions, with the clearest genotype-associated differences observed in G3. Interestingly, differences remained detectable in G5 across several treatment lineages, but they were not significant. These results suggested that the absence of GSNOR primarily affected regional methylation patterns and resulted in measurable changes in genome-wide methylation levels.

WT treatments exhibited ∼1k-∼3k DMRs across all three sequence contexts. By contrast, tens of thousands of DMRs were detected in *gsnor-ko*, with the greatest absolute numbers occurring in the CG context (∼30K). A large number of DMRs (from ∼1k in CHG to 25k in CG) were already present in the comparison between *gsnor-ko* and WT controls, demonstrating substantial baseline methylome divergence between the genotypes. Most environmental treatments further increased the regional methylation differences in *gsnor-ko*. The direction of methylation change also differed strongly between the genotypes. Although both hyper- and hypomethylated DMRs were detected in *gsnor-ko*, hypermethylated regions predominated across treatments and generations (Figure 1; Supplemental Figure S4; Supplemental File S2). The largest absolute hyper-DMR burden occurred in the CG context. In contrast, the strongest relative bias toward hypermethylation was observed in CHG and CHH, where comparatively few hypo-DMRs (∼1k) were detected.

**Figure 1.**
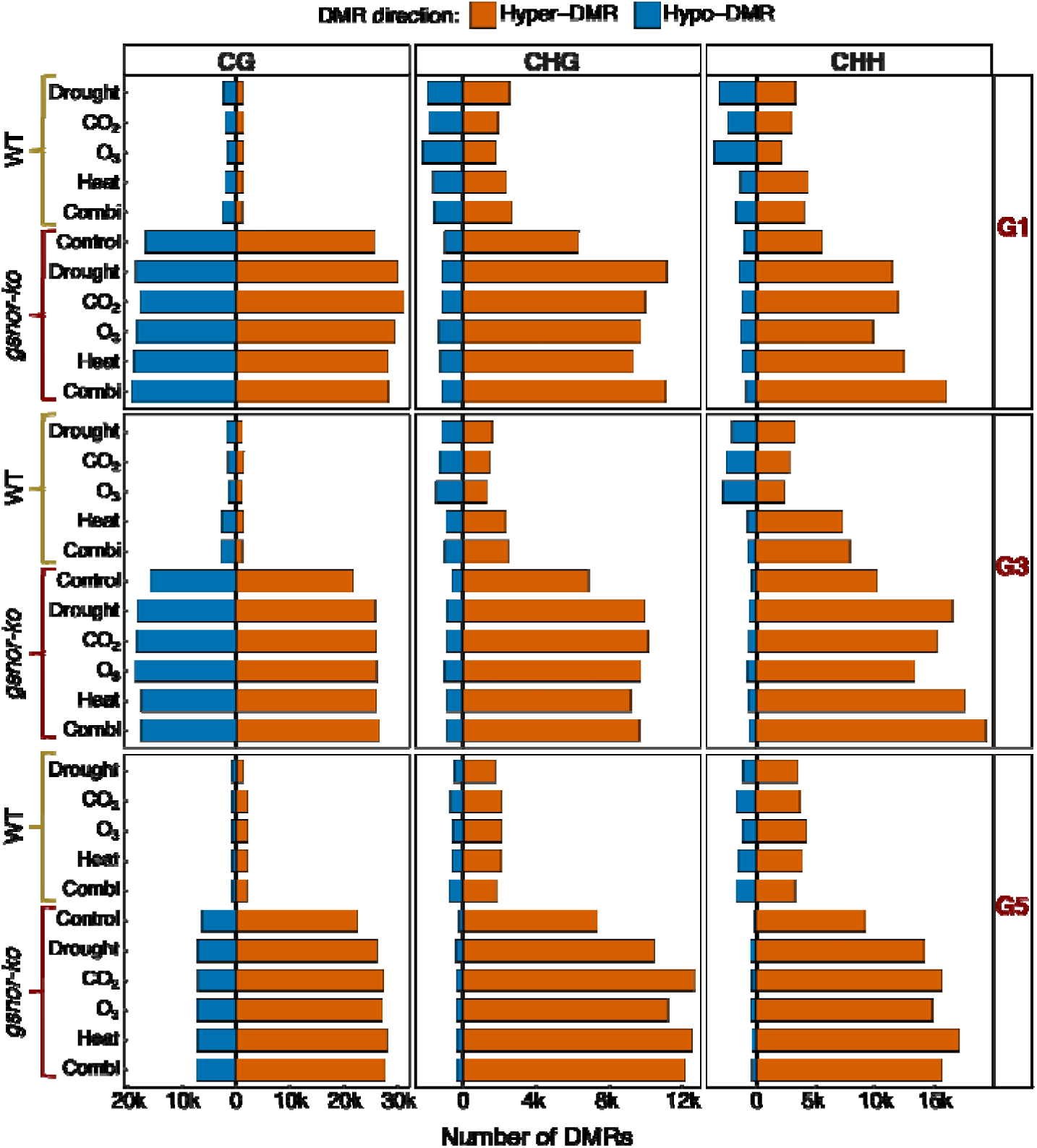
Differential methylation across generations in wild type and GSNOR-deficient *A. thaliana*. Numbers of hypermethylated (orange) and hypomethylated (blue) differentially methylated regions (DMRs) in wild type (WT) and *gsnor-ko* plants in G1, G3 and G5, separated by CG, CHG and CHH sequence context. All comparisons used WT Control from the corresponding generation as the reference. Treatments were applied during G1–G3, followed by two generations under control conditions; G5 labels therefore indicate ancestral treatment lineages. DMRs were identified using jDMRgrid with 200-bp windows, a 50-bp step, a minimum of 10 cytosines per context, exclusion of intermediate methylation calls and replicate.consensus = 1. Hyper- and hypomethylated regions were extracted from the filtered state-call matrices. WGBS included three biological replicates per control group and two per treatment group in each sampled generation. Bars represent region counts rather than replicate means; leftward bars indicate hypomethylated DMR counts.

Consistent with these patterns, the hyper-hypo DMR ratio was consistently greater in *gsnor-ko* than in WT (Figure 2; Supplemental File S2). In *gsnor-ko*, the ratio remained above 1 across the conditions examined, indicating an overall predominance of hypermethylation. In WT, CG hyper:hypo ratios were commonly below or near 1 in G1 and G3, indicating similar numbers of hyper- and hypo-DMRs or a modest predominance of hypomethylation. The CG ratio subsequently increased in several WT treatment lineages by G5. WT CHG and CHH ratios were generally lower than those of *gsnor-ko* and showed more treatment-dependent trajectories. For example, CHH ratios increased at G3 under heat and combination treatments but declined again by G5. Therefore, WT showed context- and treatment-dependent changes during recovery, whereas the methylation response of *gsnor-ko* was characterised by a stronger and more persistent hypermethylation bias.

**Figure 2.**
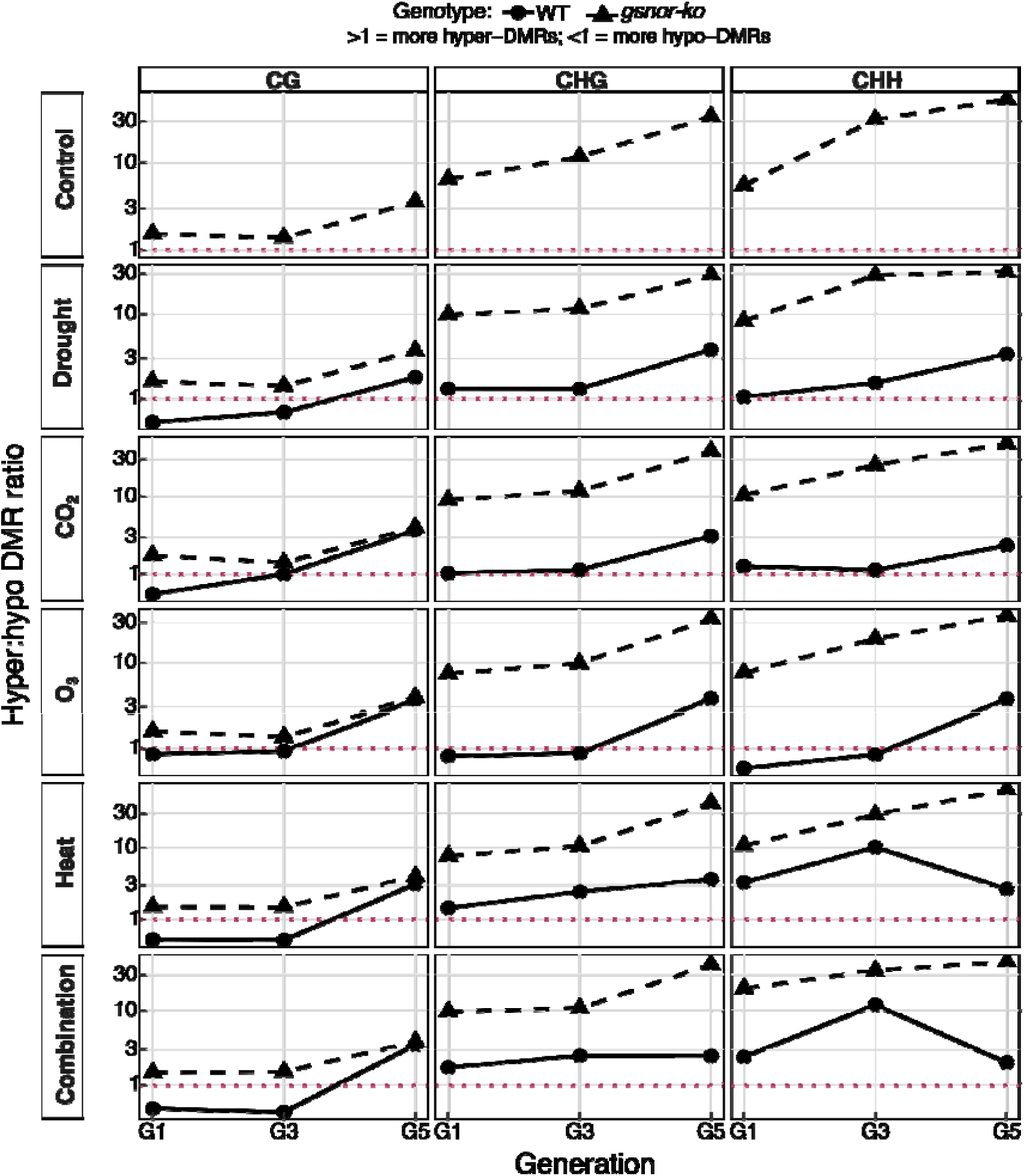
Changes in the balance between hypermethylated and hypomethylated regions across generations. Hyper DMR ratios in WT and *gsnor-ko* plants across G1, G3 and G5, separated by treatment lineage and methylation context. Ratios were calculated by dividing hypermethylated by hypomethylated DMR counts, using WT Control from the corresponding generation as the reference. Circles connected by solid lines represent WT, and triangles connected by dashed lines represent *gsnor-ko*. The dotted magenta line marks a ratio of 1, indicating equal numbers of hypermethylated and hypomethylated DMRs. Values above and below 1 indicate a predominance of hypermethylation and hypomethylation, respectively. The vertical axis is logarithmic. G5 labels indicate ancestral treatments followed by two generations under control conditions. Ratios are descriptive summaries of the DMR counts shown in Figure 1.

Although *gsnor-ko* Control already exhibited substantial baseline methylome divergence from WT, the environmental-treatment lineages generally contained more DMRs (∼1.5-2k) than the mutant control comparison, indicating additional treatment-associated remodelling superimposed on the genotype-associated background (Figure 1). Hyper-DMRs shared across G1, G3 and G5 represented 39.9–42.8% of CG, 8.4–9.4% of CHG and 3.1–3.4% of CHH DMR unions in *gsnor-ko*, compared with 0–0.7%, 0–0.1% and 0–0.1%, respectively, in WT (Supplemental Figures S5–S7). Generation-specific DMRs were also detected in both genotypes, indicating that cross-generation retention occurred alongside continued methylome remodelling. Similarly, the hyper-DMR core shared among the displayed treatment sets represented 34.6–47.5% of CG, 14.0–18.0% of CHG and 8.2–9.2% of CHH DMR unions in *gsnor-ko*, compared with 0.5–0.9%, 0.7–1.2% and 0.9–1.5%, respectively, in WT (Supplemental Figures S8–S10). These intersections indicate a large shared mutant component together with additional treatment- and generation-specific changes

DMR localisation remained strongly context dependent: CG DMRs were most frequently associated with gene bodies, whereas CHG and CHH DMRs occurred predominantly in TE-associated regions (Supplemental Figures S11–S13). Although this broad context-specific pattern was conserved, the relative representation of gene-body- and TE-associated DMRs differed quantitatively between genotypes. Overall, loss of GSNOR was associated with baseline methylome divergence, an additional treatment-associated DMR burden, a pronounced hypermethylation bias and reduced methylome resetting across generations.

### 3.2 Methylation divergence and DMR persistence differ between WT and *gsnor-ko*

We next used AlphaBeta analysis to determine whether methylation divergence accumulated differently between genotypes and treatments. In the CG context, fitted divergence trajectories were consistently higher in *gsnor-ko* than in WT (Figure 3A). Nominally significant treatment effects were detected within both genotypes, while comparisons between genotypes identified differences under control, CO□, O□, and heat conditions (Supplemental Table S1). CHG methylation dynamics also differed among selected treatments and genotypes, with the strongest genotype difference occurring under the combination treatment. By contrast, no AlphaBeta comparison was significant in the CHH context (Supplemental Figure S14; Supplemental Table S1; Supplemental File S3). Thus, the loss of GSNOR predominantly affected the dynamics of CG- and CHG-methylation divergence. Full parameter estimates and *P* values are provided in Supplemental Table S1.

**Figure 3.**
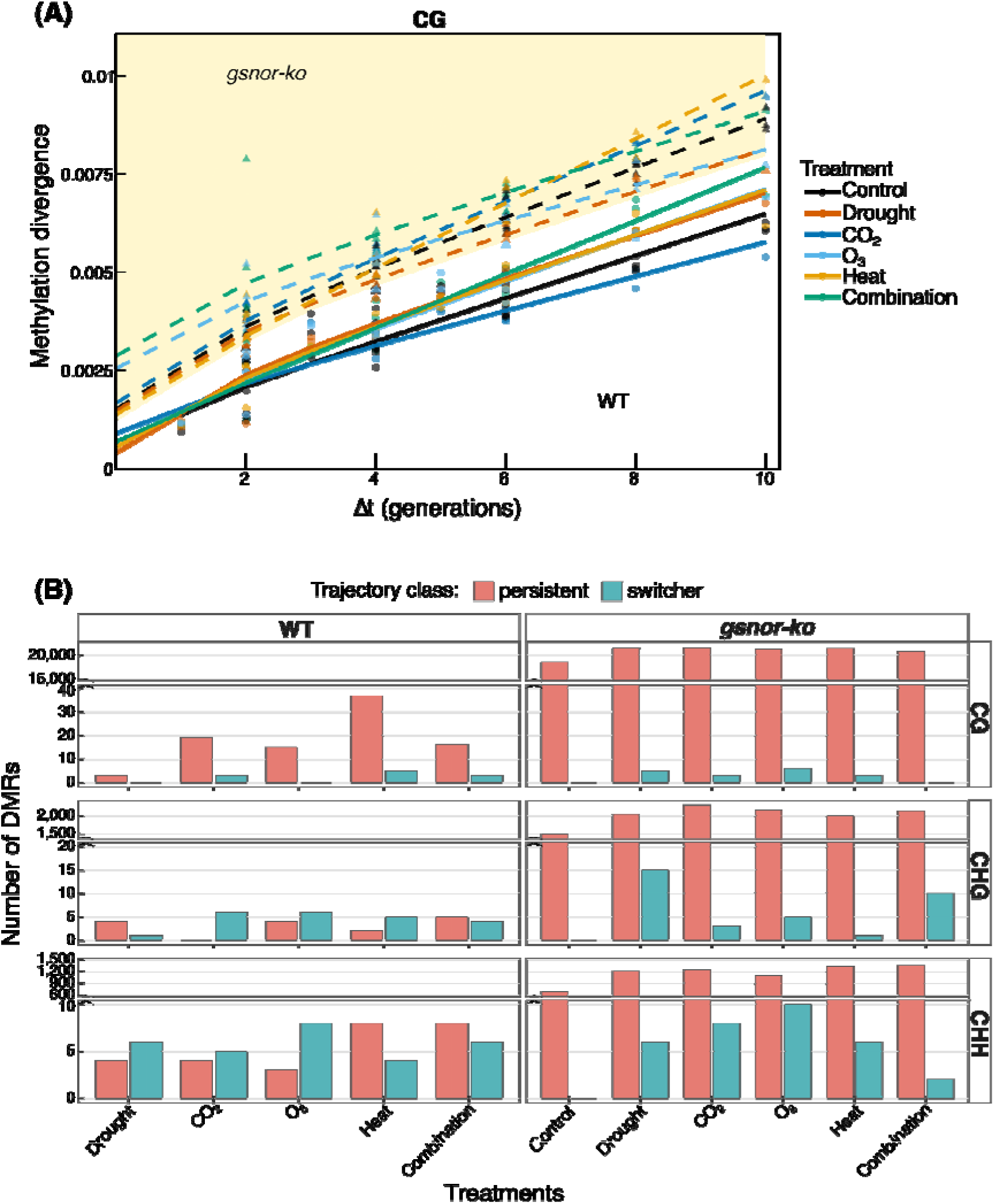
CG methylation divergence and persistence of regional methylation differences across generations. **(A)** Observed CG methylation divergence and fitted AlphaBeta trajectories for WT and *gsnor-ko* treatment lineages. Circles and solid lines represent WT observations and fitted trajectories, respectively; triangles and dashed lines represent *gsnor-ko*. Colours distinguish treatments. Δt denotes the summed generational distance separating two sampled individuals through their common ancestor. Methylationgain (α) and methylation-loss (β) parameters were estimated using the AlphaBeta ABneutral model, with a posteriorMaxFilter setting of 0.99 and 50 optimisation starts. Parameter uncertainty was assessed using 100 bootstrap iterations. Pale-yellow shading highlights the area above a boundary placed just below the lowest fitted *gsnor-ko* trajectory at each Δt; it is a visual aid and does not represent a confidence interval or significance threshold. **(B)** Numbers of persistent (salmon) and switcher (turquoise) DMRs by genotype, treatment lineage and methylation context. All differential methylation states were defined relative to WT Control within the corresponding generation. Persistent DMRs retained the same direction at G1, G3 and G5; switcher DMRs were detected at all three sampling points but changed direction at least once. Vertical-axis breaks accommodate differences in abundance. Bars show region counts rather than replicate means.

We subsequently classified DMRs according to their G1–G3–G5 trajectories. A persistent DMR was defined as a region that remained differentially methylated in the same direction in all three generations, either hyper–hyper–hyper or hypo–hypo–hypo. A switcher was present as a DMR in all three generations but changed direction at least once. To distinguish treatment-associated persistence from the baseline effect of the mutation, WT treatments and *gsnor-ko* control and treatments were compared with WT Control at each generation. WT contained 3–37 persistent CG DMRs across the five treatments, compared with 17–74 in *gsnor-ko* (Figure 3B; Supplemental File S4). Persistent CHG DMRs ranged from 0–5 in WT and 1–12 in *gsnor-ko*, while persistent CHH DMRs ranged from 3–8 in WT and 0–5 in *gsnor-ko*. By contrast, the separate baseline comparison of *gsnor-ko* Control with WT Control identified 18,593 persistent CG, 1,478 CHG, and 693 CHH DMRs. Thus, the very large persistent DMR population in *gsnor-ko* primarily represented a stable baseline genotype difference rather than treatment-specific inheritance.

Because the initial number of treatment-associated DMRs differed among genotypes, treatments, and sequence contexts, persistence was also expressed as the percentage of G1 DMRs that retained the same methylation direction in G3 and G5. In the CG context, 0.9–4.5% of G1 DMRs persisted in *gsnor-ko*, compared with 0.1–1.2% in WT. Persistence was substantially lower in the non-CG contexts, reaching a maximum of 0.6% for CHG DMRs in *gsnor-ko* and approximately 0.1% for CHG DMRs in WT and for CHH DMRs in both genotypes. Therefore, *gsnor-ko* showed greater treatment-specific retention mainly in the CG context, but most treatment-associated methylation changes were not maintained across all three generations. These results do not indicate a general loss of methylome flexibility in *gsnor-ko*; instead, they distinguish a highly persistent baseline genotype signature from a smaller increase in the retention of treatment-associated CG DMRs.

Switcher DMRs were fewer than persistent DMRs in *gsnor-ko*. Depending on treatment and context, only 0–15 DMRs remained present across all three generations. WT displayed a few switcher DMRs, although in CHG and CHH their numbers occasionally equaled or exceeded the small number of persistent regions. The alluvial plots illustrated these trajectories and showed that same-direction retention represented a major component of the *gsnor-ko* DMR population, whereas flows between hypermethylated, hypomethylated, and absent states accounted for the remaining dynamic regions (Supplemental Figure S16A-C; Supplemental File S4).

The genomic distribution of persistent DMRs was context-dependent (Supplemental Figures S17 and S18). Persistent CG DMRs most frequently overlapped gene bodies, whereas persistent CHG and CHH DMRs were primarily associated with TEs. Promoter-associated and intergenic DMRs constituted smaller fractions of the persistent sets. Switcher DMRs showed more variable annotation profiles, consistent with their low abundance, but CHG and CHH switchers were also frequently associated with TEs (Supplemental Figures S17 and S19). These annotation patterns were broadly similar between genotypes, indicating that the major difference between WT and *gsnor-ko* was the extent of DMR retention rather than a complete redistribution among genomic features.

### 3.3 Persistent and switching DMRs highlight genes associated with stress and epigenetic regulation

Candidate genes associated with persistent DMRs were organised into functional groups related to epigenetic and small-RNA regulation, DNA methylation and chromatin organisation, redox and stress signalling, hormone, calcium and nutrient signalling, and reproduction or transgenerational regulation (Figure 4A; Supplemental File S5). Persistent promoter- and gene-body DMRs occurred across all these functional classes and included both hypermethylated and hypomethylated regions. These associations were detected predominantly in *gsnor-ko*, frequently across multiple treatments, whereas only a limited number of WT-specific persistent DMRs were identified. Their retention in the same methylation direction across G1, G3, and G5 indicates that methylation changes affecting genes involved in stress responses and epigenetic regulation remained detectable after treatment withdrawal.

**Figure 4.**
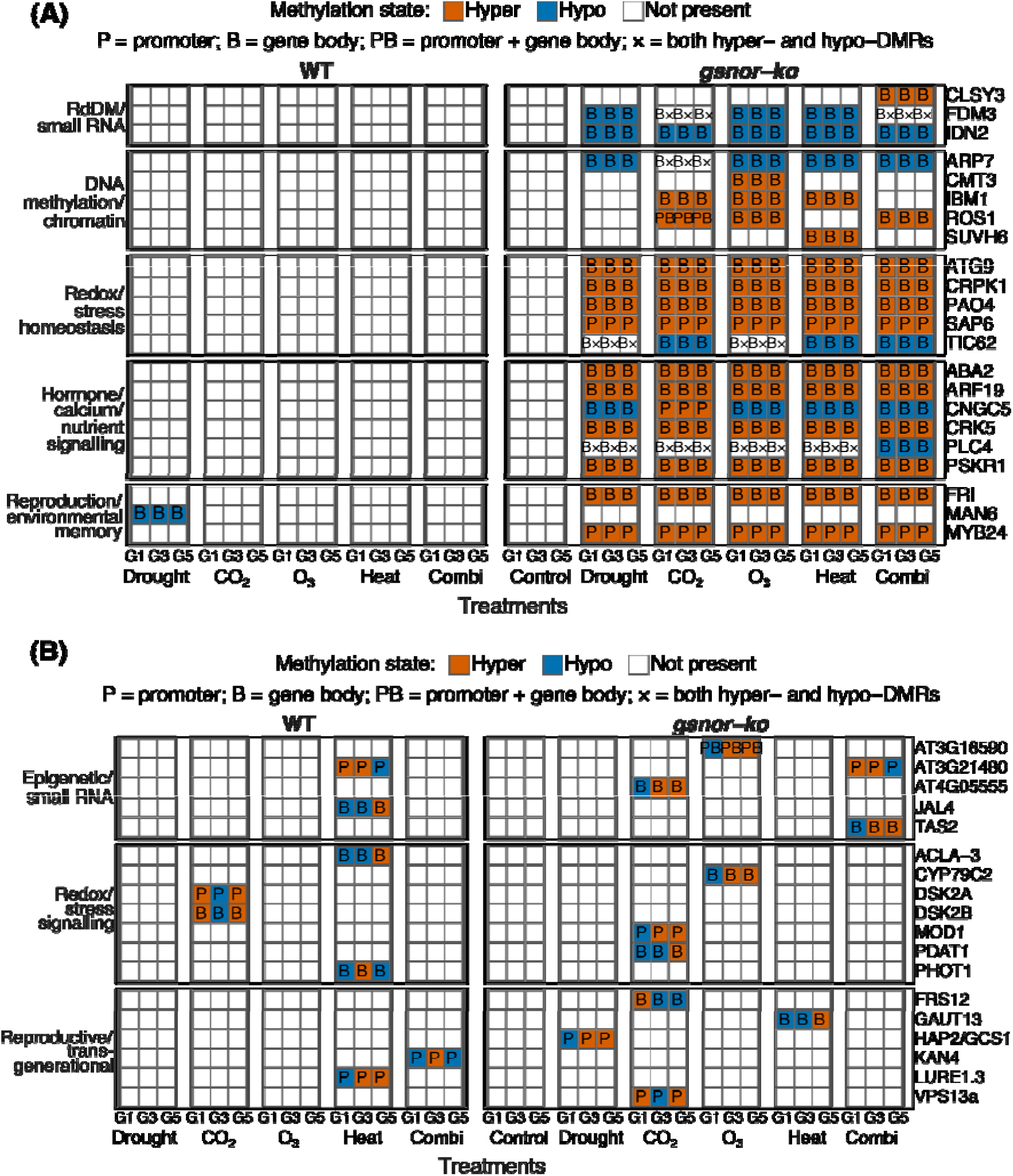
Persistent and switching methylation patterns at selected candidate genes. Heatmaps showing candidate genes associated with **(A)** persistent and **(B)** switcher DMRs in WT and *gsnor-ko* plants. Methylation states were defined relative to WT Control within the corresponding generation. Columns within each treatment lineage represent G1, G3 and G5. Orange and blue indicate hypermethylated and hypomethylated DMR associations, respectively. White cells indicate no displayed DMR association, except where × denotes both hypermethylated and hypomethylated DMRs associated with the same gene. P, B and PB indicate promoter, gene-body, and combined promoter/gene-body associations, respectively. Promoters were defined as 1 kb upstream of the annotated transcription start site. Persistent DMRs retained the same direction across all three sampled generations; switcher DMRs changed direction while remaining detectable at all three sampling points. Genes were selected and grouped according to literature-supported functions; these categories do not represent statistical enrichment results. G5 labels indicate ancestral treatment lineages following two generations under control conditions.

Switcher DMRs were associated with genes belonging to functional classes related to epigenetic and small-RNA regulation, redox and stress signalling, hormone, calcium and nutrient signalling, and reproduction or transgenerational regulation. These patterns occurred in both genotypes, although switcher-associated candidates were more widely distributed across treatments in *gsnor-ko* (Figure 4B; Supplemental Figure S20; Supplemental File S5).

### 3.4 DMRs correlating with altered gene expression

We next performed transcriptome-guided cross-omics validation using the same epigenetic-regulator genes previously identified in our RNA-seq study, which showed transgenerational expression differences between WT and *gsnor-ko* (Behl et al., 2026). Gene-associated DMRs were detected for many of these independently defined RNA-seq candidates. WT generally displayed fewer and more treatment- or generation-restricted methylation changes. By contrast, *gsnor-ko* showed a denser distribution of promoter- and gene-body DMRs across treatments and generations, including hypermethylated, hypomethylated, and mixed methylation states. Particularly extensive methylation differences were observed among genes associated with DNA methylation and silencing, Polycomb regulation, histone and chromatin organisation, RNA methylation, and methyl-donor metabolism (Supplemental Figure S21). This demonstrates that genes differing transgenerationally between WT and *gsnor-ko* at the transcript level are frequently associated with local methylation differences in the corresponding multigenerational WGBS dataset. This convergence provides cross-omics support for an association between GSNOR-dependent transcriptomic reconfiguration and altered methylation at epigenetic regulatory genes. However, because the analysis is correlative, it does not establish that the observed DMRs directly caused the corresponding expression differences.

To further determine whether methylation changes at the selected candidate genes coincided with transcriptional responses, the WGBS results were integrated with RNA-seq data. Methylation and expression measurements were matched by genotype, treatment, and generation. Integration of persistent DMRs with RNA-seq responses revealed marked differences between WT and *gsnor-ko* (Figure 5A; Supplemental File S6). In WT, candidates generally displayed fewer significant expression responses, consistent with the comparatively limited occurrence of persistent candidate-associated DMRs. By contrast, a larger proportion of the persistent candidates in *gsnor-ko* showed altered expression in one or more treatment–generation combinations. These expression responses were observed at genes carrying persistent promoter DMRs, gene-body DMRs, or DMRs overlapping both genomic features. Several candidates showed expression responses across multiple generations or treatments, whereas others exhibited expression changes restricted to a particular environmental condition or generation. Switcher candidates also displayed genotype- and condition-dependent transcriptional responses (Figure 5B; Supplemental File S6). Significant expression changes were detected for a subset of switcher genes, particularly in *gsnor-ko*, although the responses were generally more heterogeneous than those associated with persistent DMRs. Changes in methylation direction across G1, G3, and G5 were not invariably accompanied by corresponding reversals in gene-expression direction. Instead, some switcher candidates exhibited generation-specific expression responses, while others retained similar expression directions despite transitions between hypermethylated and hypomethylated states.

**Figure 5.**
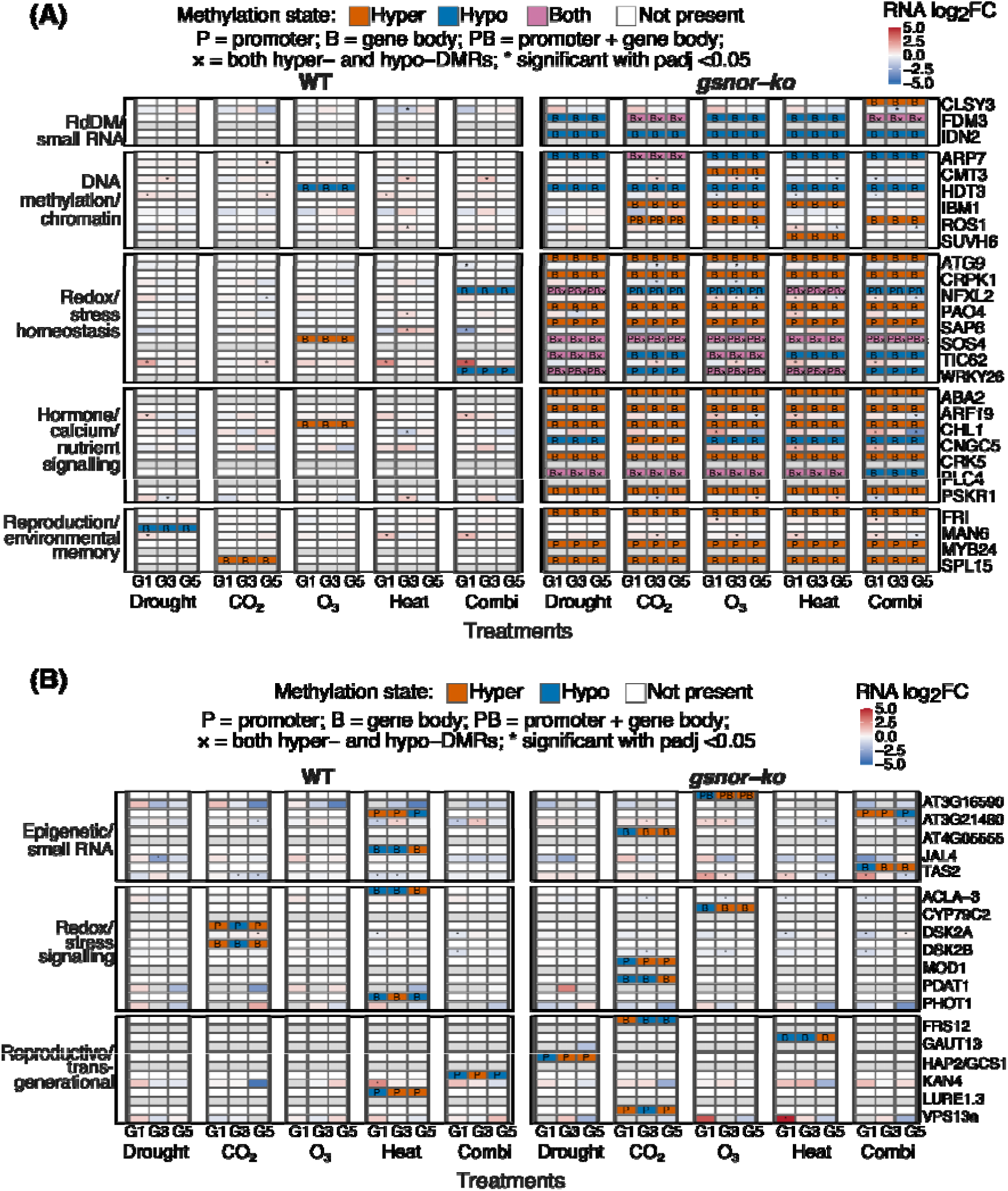
Integration of candidate-gene methylation patterns with transcriptional responses. Paired methylation and expression heatmaps for genes associated with **(A)** persistent and **(B)** switcher DMRs in WT and gsnor-ko plants. Columns within each treatment lineage represent G1, G3 and G5. For each gene, the upper track displays DMR associations and the lower track displays the corresponding RNA-seq log fold change from Behl et al. (2026). Methylation states were defined relative to WT Control within the corresponding generation. Orange, blue and magenta indicate hypermethylated, hypomethylated and combined hyper-/hypomethylated associations, respectively; white indicates no displayed DMR association. P, B and PB denote promoter, gene-body, and combined promoter/gene-body associations; × denotes both methylation directions. RNA colours represent unshrunken log fold changes, including nonsignificant results, with blue indicating negative and red indicating positive values. Asterisks indicate DESeq2-adjusted P < 0.05, without an additional fold-change cutoff. Grey RNA cells indicate unavailable matched expression values; near-white expression colours indicate values close to zero. WGBS and RNA-seq results were matched by genotype, treatment and generation. G5 labels indicate ancestral treatment lineages following two generations under control conditions.

### 3.5 Persistent differential TE methylation is associated with altered expression of nearby genes

Persistent DMRs within TEs were examined separately from gene-associated DMRs to determine whether stable TE methylation changes coincided with transcriptional responses in neighbouring genes. TEs were classified according to their annotated TE class or superfamily, including DNA/MuDR, RC/Helitron, LTR/Copia, and LTR/Gypsy elements (Figure 6; Supplemental File S7). Persistent TEs (DMRs) were detected predominantly in *gsnor-ko*, whereas corresponding persistent TE methylation was limited or absent in WT for most candidates. In *gsnor-ko*, several TE loci retained the same hypermethylated or hypomethylated state across G1, G3, and G5 under one or more treatment conditions. The distribution of these persistent states varied across TE classes and treatments, indicating that TE methylation retention was locus- and condition-dependent rather than uniform. Several genes located near persistently methylated TEs showed altered expression, with the majority of these responses occurring in *gsnor-ko*. However, significant expression differences were not detected for every persistently methylated TE–gene pair. The direction of nearby-gene expression did not show a uniform relationship with the methylation state of the associated TE. Persistent TE hypermethylation coincided with reduced expression for some nearby genes but with increased, unchanged, or statistically nonsignificant expression for others. Expression responses also varied among treatments and generations, even when the corresponding TE retained the same methylation direction. These patterns suggest that the relationship between TE methylation and nearby-gene expression depends on the individual TE–gene pair and its genomic and environmental context.

**Figure 6.**
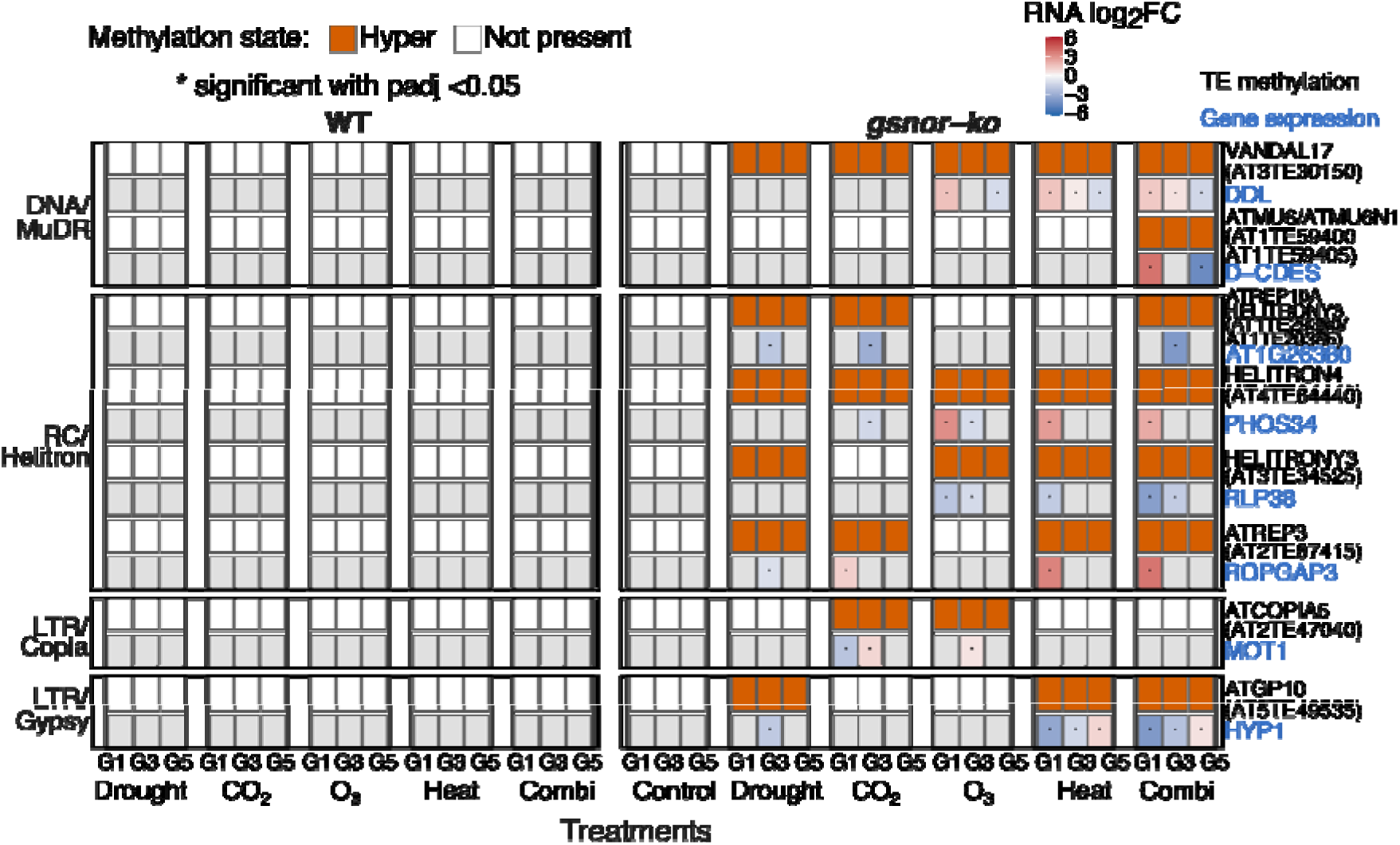
Persistent TE-associated hypermethylation and expression of neighbouring genes. Paired heatmaps showing eight selected transposable-element (TE)–gene candidate groups in WT and *gsnor-ko* plants, organised by TE superfamily: DNA/MuDR, RC/Helitron, LTR/Copia and LTR/Gypsy. The upper track displays persistent TE-associated methylation and the lower track displays the neighbouring gene’s RNA-seq log fold change. Columns within each treatment lineage represent G1, G3 and G5. Orange indicates persistent hypermethylation relative to WT Control within the corresponding generation; white indicates no displayed persistent TE-associated DMR. Persistent DMRs retained the same differential methylation direction across all three sampled generations. Candidate TEs were linked to nearby genes using a 1.2-kb proximity threshold. RNA results from Behl et al. (2026) were retained only for significant, condition-matched TE–gene associations with DESeq2-adjusted P < 0.05, without an additional fold-change cutoff. Blue and red indicate negative and positive log fold changes, respectively; asterisks mark significant expression results. Grey RNA cells indicate absence from the filtered, condition-matched significant TE–gene table and do not establish unchanged expression. G5 labels indicate ancestral treatment lineages following two generations under control conditions.

## 4. DISCUSSION

This study shows that GSNOR-dependent nitric oxide homeostasis is associated with both the magnitude and the persistence of DNA methylation variation during repeated environmental exposure and subsequent recovery. The main findings are that loss of GSNOR produced a strongly divergent baseline methylome, greatly increased the number of DMRs, favoured hypermethylation, and reduced the resetting of regional methylation states after treatment withdrawal.

### 4.1 Loss of GSNOR shifts the methylome towards regional hypermethylation

The separation of WT and *gsnor-ko* control methylomes, along with the large number of DMRs already present under control conditions, demonstrates that GSNOR has a substantial genotype-dependent effect on the methylome. This finding is consistent with earlier evidence that GSNOR1 contributes to chromatin accessibility, DNA demethylation, and the expression of TEs and stress-responsive genes (Rudolf et al., 2021). *Rudolf et al.* linked GSNOR1 activity to the methylation index and to H3K9me2 abundance, providing a mechanistic precedent for altered methylation in *gsnor-ko*. Other studies have shown that nitric oxide regulates histone acetylation through GSNOR- and HDA6-dependent processes and that S-nitrosylation can directly modify the activity and activities of HDACs (Ageeva-Kieferle et al., 2021; Mengel et al., 2017; Zheng et al., 2023). These studies established a connection between nitric oxide homeostasis and chromatin regulation, but they did not determine how loss of GSNOR affects DNA methylation during repeated stress and recovery across generations.

Our results extend this literature in three respects. First, the effect of GSNOR loss was not restricted to a particular stress: a high DMR burden was detected in the mutant control and across drought, elevated CO_2_, O_3_, heat, and combined-treatment lineages. Second, differences in average methylation were comparatively modest, whereas tens of thousands of DMRs were detected in *gsnor-ko*. This distinction is important because opposing or spatially restricted methylation changes can produce limited changes in the genome-wide mean while extensively remodelling regulatory regions. Third, *gsnor-ko* showed a persistent bias towards hypermethylation, especially as a high absolute number of CG hyper-DMRs and a marked hyper: hypo imbalance in CHG and CHH. These results demonstrate that loss of GSNOR alters the balance of methylation states but do not establish whether this phenotype results from reduced demethylation, increased methylation establishment or maintenance, or another indirect mechanism, as also reported by (Rudolf et al., 2021).

The genomic distribution of the DMRs followed the established architecture of the Arabidopsis methylome. CG DMRs were frequently associated with gene bodies, whereas CHG and CHH DMRs were concentrated at TEs. CG, CHG, and CHH methylation are maintained by partly distinct pathways, and non-CG methylation is particularly important for heterochromatin and TE silencing (Johannes & Schmitz, 2019; Law & Jacobsen, 2010; Stroud et al., 2014; Zhang et al., 2018). Although the proportions of gene- and TE-associated DMRs differed between WT and *gsnor-ko*, both genotypes showed the same general context-dependent pattern. Thus, loss of GSNOR mainly affected the number and direction of DMRs rather than producing a completely different distribution among genes, TEs, promoters, and intergenic regions.

### 4.2 GSNOR is associated with methylation divergence and resetting after stress withdrawal

The AlphaBeta analysis and the region-level trajectory analysis provide complementary views of methylome dynamics. The consistently higher CG divergence fitted for *gsnor-ko*, together with the significant difference between the WT and mutant controls, indicates that altered methylation dynamics pre-existed the treatment-dependent responses. Significant effects of CO_2_, O_3_, and heat relative to the *gsnor-ko* control further show that environmental conditions modified this altered baseline. The absence of a significant difference between *gsnor-ko* Combination and *gsnor-ko* Control does not mean that the combined treatment had no effect on the methylome; probably, already divergent mutant control reduced the additional contrast attributable to the combined treatment. By comparison, WT Combination differed from WT Control for the methylation-gain parameter, indicating that the same environmental regime was superimposed on a less divergent baseline in WT.

Previous studies have reported conflicting evidence regarding the stability of stress-associated methylation. Hyperosmotic priming can generate locus-specific chromatin and DNA methylation states that influence later responses, but much of this memory is transient or extensively reset when stress is removed (Sani et al., 2013; Wibowo et al., 2016, 2018). Repeated drought experiments have found few conserved stress-induced DMRs and have attributed much of the apparent transgenerational variation to stochastic or pre-existing differences, particularly in repetitive regions (Ganguly et al., 2017). Other studies have likewise concluded that transgenerational stress memory is not a universal response in Arabidopsis (Pecinka et al., 2009). These findings emphasise that persistence depends on genotype, stress regime, tissue, developmental stage, and the number of untreated generations used to test recovery.

In the present study, the marked contrast between WT and *gsnor-ko* is an important finding. Only a very small fraction of WT G1 DMRs retained the same direction through G3 and G5, consistent with efficient resetting and with earlier reports of a generally stable WT methylome. In *gsnor-ko*, however, approximately 44–46% of G1 CG DMRs, 17–21% of CHG DMRs, and 8–10% of CHH DMRs remained in the same direction across G1, G3 and G5. Persistence, therefore, remained much greater in the mutant even after normalisation to its larger G1 DMR population. Because G4 and G5 were grown under control conditions, the G5 states cannot be explained by continued direct exposure. They instead represent methylation differences retained within the descendant lineage after two generations of recovery. On the other hand, switcher DMRs were rare, indicating that complete reversals between hypermethylated and hypomethylated states were not the dominant outcome. The combination of extensive same-direction retention and a smaller set of directional switches suggests that loss of GSNOR changes both the capacity to erase methylation differences and the route by which some loci are remodelled during recovery. This provides a genome-wide extension of the known role of nitric oxide as a regulator of chromatin state and supports a model in which balanced S-nitrosothiol signalling contributes to epigenomic homeostasis (Ageeva-Kieferle et al., 2021; Lindermayr et al., 2020).

### 4.3 Persistent and switching DMRs implicate interconnected regulatory pathways

We identified genes that overlapped with persistent or switching DMRs and grouped them by their known biological functions. These groups included small-RNA pathways, DNA methylation and chromatin organisation, redox and stress signalling, hormone responses, and plant development. This literature-based classification was used to identify biologically relevant candidates and should not be interpreted as a statistical enrichment analysis.

Persistent DMRs associated with *CLSY3, FDM3, and IDN2* are notable because these genes function in or are connected to RNA-directed DNA methylation, while *CMT3, IBM1, ROS1*, and *SUVH6* participate in the establishment, interpretation, or removal of repressive methylation and chromatin states. The presence of persistent promoter- or gene-body DMRs at these loci suggests feedback between altered methylation and the machinery that regulates it. However, a DMR associated with a methylation regulator does not by itself demonstrate altered protein activity or pathway output, and this interpretation must remain a hypothesis until tested experimentally. The persistent candidates also connected methylation dynamics with pathways expected to be sensitive to altered nitric oxide homeostasis. Genes related to redox balance, autophagy, photosynthetic electron transfer, ABA and auxin signalling, calcium signalling, and reproductive development carried persistent DMRs predominantly in *gsnor-ko*. GSNOR is already known to influence development, fertility, thermotolerance, auxin signalling, and stress responses (Feechan et al., 2005; Lee et al., 2008; Shi et al., 2015). Our results do not show that methylation causes these phenotypes, but they identify stable local epigenomic states at genes positioned within the same biological processes.

Switcher DMRs provided a complementary set of candidates whose methylation direction changed across generations. These candidates were associated mainly with epigenetic regulation, stress responses, and plant development, suggesting that some loci continued to be remodelled during the transition from repeated treatment to recovery. As with the persistent DMRs, these functional associations are descriptive and require experimental validation.

### 4.4 Methylation–expression relationships are locus- and context-dependent

Integration of the WGBS and RNA-seq datasets showed that more persistent and switcher candidates exhibited altered expression in *gsnor-ko* compared with WT. Because methylation and expression were matched by genotype, treatment, and generation, these analyses resolve associations that would be obscured by comparisons across unmatched conditions. Persistent DMRs were often accompanied by expression changes in one or more treatment–generation combinations, whereas switcher candidates tended to show more heterogeneous responses. The broader transcriptional response in *gsnor-ko* is consistent with our previous finding that loss of GSNOR produced stronger persistence or reconfiguration of stress-associated expression states during recovery (Behl et al., 2026). This convergence supports an association between GSNOR-dependent transcriptome reconfiguration and local methylation differences at epigenetic regulators. It does not establish whether methylation preceded expression change, resulted from altered transcription or chromatin state, or participated in a reciprocal feedback loop. There was, however, no universal inverse relationship between methylation and expression. Promoter hypermethylation can restrict transcription in some settings, but gene-body methylation has more variable associations with transcription, and the effect of a DMR depends on its position relative to regulatory elements, its sequence context, the surrounding chromatin, and the cell types contributing to the bulk tissue (Law & Jacobsen, 2010; Zhang et al., 2018). Consistent with this complexity, hypermethylation in our dataset was associated with decreased, increased, unchanged, or non-significant expression, depending on the locus and condition; hypomethylation was similarly not synonymous with activation. Ganguly et al. (2017) likewise found that drought-associated methylation variants did not show a general relationship with drought-responsive expression. Conversely, Wibowo et al. (2016) demonstrated at the CNI1 locus that a stress-induced DMR can regulate a long non-coding RNA and influence gene expression, illustrating that causal methylation–expression coupling may exist at selected loci.

Accordingly, the value of the integrated heatmaps is not evidence for a single direction of regulation but the identification of candidates with coordinated, condition-matched molecular changes. Loci that combine a persistent promoter DMR with reproducible expression differences across generations are especially suitable for targeted validation. Bisulfite amplicon sequencing, locus-specific expression assays, chromatin profiling, and targeted methylation or demethylation would be required to distinguish causal regulation from correlated responses to a common upstream change in nitric oxide or chromatin state.

### 4.5 Persistent TE methylation extends the connection between GSNOR and transposon regulation

The number of persistent CHG and CHH DMRs at TEs is consistent with the central role of non-CG methylation in heterochromatic TE silencing (Liang et al., 2022; Stroud et al., 2014). It also agrees with the independent observation that GSNOR1 contributes to TE demethylation and expression (Rudolf et al., 2021). In the present study, persistent TE-associated DMRs occurred mainly in *gsnor-ko* and included DNA/MuDR, RC/Helitron, LTR/Copia, and LTR/Gypsy elements. Their persistence across G1, G3, and G5 extends the earlier single-generation connection between GSNOR and TEs by showing that altered TE methylation can remain detectable through repeated exposure and subsequent recovery. Some genes located near persistently methylated TEs also showed altered expression, but the direction of expression could not be predicted from TE methylation alone. We tested this because TEs can influence nearby genes through local chromatin spreading, regulatory sequences, or TE-derived small RNAs, whereas many methylated TEs remain transcriptionally silent and have no measurable effects on adjacent genes (Ahmed et al., 2011; Chuong et al., 2017; Dubin et al., 2018; Ito, 2022; Ramakrishnan et al., 2021). The paired TE–gene analysis therefore identifies potential regulatory neighbourhoods rather than proving direct cis regulation. Moreover, this study did not measure TE mobilisation, and altered TE methylation or expression should not be equated with transposition.

### 4.6 Independent comparisons support a reproducible subset of GSNOR-associated DMRs

Comparison with published *Arabidopsis* stress-methylome datasets showed that some of the persistent and switching DMRs, candidate genes, and TEs identified in the present study were also detected in independent experiments (Supplemental Figures S22–S24; Supplemental File S8) (Behl et al., 2025). CHH showed the largest absolute number of overlapping DMRs, but CHH DMRs were also the most abundant overall. Therefore, this result does not mean that CHH changes were proportionally more reproducible than CG or CHG changes. Variation among studies was expected because they differed in stress conditions, plant age, tissue, genotype, sequencing depth, and methods used to identify DMRs.

The published *gsnor1-3*-versus-WT dataset provided a more direct comparison of the baseline mutant phenotype (Supplemental Figure S25; Supplemental File S9). Of the 292 published DMRs, 215 (73.6%) overlapped at least one DMR detected in the current G1 *gsnor-ko* Control comparison, and 211 of these regions showed the same methylation direction. Viewed from the opposite direction, 1,378 of the 55,942 current G1 DMRs (2.5%) overlapped a published region. These percentages differ because the published DMR set was small and focused, whereas the present analysis detected many more context-specific and overlapping 200-bp windows. One published region could therefore overlap several current DMR windows. The high agreement in methylation direction among overlapping regions supports the presence of a reproducible subset of GSNOR-associated DMRs. However, the published dataset did not classify individual DMRs as CG, CHG, or CHH, so concordance within the different sequence contexts could not be assessed.

The independently supported G1 regions were subsequently followed across G3 and G5 to determine whether their methylation states persisted or changed over generations (Supplemental Figure S25; Supplemental File S9). We also distinguished DMRs shared with *gsnor-ko* Control from those detected only in the treatment comparisons. This separated stable baseline genotype effects from additional treatment-associated responses. The analysis therefore extends the Rudolf et al. (2021) dataset by showing not only which GSNOR-associated DMRs were reproduced at G1, but also how these regions behaved during multigenerational treatment and recovery.

Future studies should validate the highest-priority regions using targeted bisulfite sequencing and determine whether their methylation states affect gene expression. Analyses of chromatin accessibility, histone modifications, small RNAs, and S-nitrosylated proteins could help explain how GSNOR-dependent redox signalling is connected to DNA methylation. For TE-associated DMRs, direct measurements of TE expression and mobilisation will be necessary, as methylation changes alone do not indicate altered TE activity.

## Supporting information

Behl et al_Supplemental Figures

Behl et al_Supplementary_Files_S1-S9

## ACKNOWLEDGEMENTS

This work (for RB and CL) was supported by the Initiative and Networking Fund of the Helmholtz Association under the call and by EpiCrossBorders: International Helmholtz-Edinburgh Research School for Epigenetics.

We thank Peiyuan Zhu, Moritz Popp, Ina Zimmer, Petra Seibel, Franz Buegger, Mustafa Oezden, Alexandros Sigalas, Georg Gerl, Ulrich Junghans, Baris Weber, and Armin Richter for their technical support during experimental work, and Zhilin Zhang and Rohan Bhardwaj for help with WGBS data preprocessing. We would also like to thank Prof. Dr. Aloys Schepers for his fruitful input during thesis meetings.

During the preparation of this work, the author(s) used AI tools (such as ChatGPT 5.64 and Grammarly 1.162.2.0) to polish the writing framework and to search for relevant research articles (literature overview). After using this tool/service, the author(s) reviewed and edited the content as needed and take(s) full responsibility for the content of the published article.

## CONFLICT OF INTEREST

Not declared

## AUTHOR CONTRIBUTIONS

Experiments were designed by CL & FJ; experimental work was conducted by RB, AG, EM, LG, AA, JPS & CL. WGBS library preparation was performed by BV. Analysis was performed by RB; statistical analysis and WGBS analysis support were provided by JPS, FJ & CL; data interpretation was performed by RB, JPS, FJ & CL; and the paper was written by RB with input from JPS, AG, FJ & CL. The written paper was reviewed by all the co-authors.

## SUPPLEMENTAL MATERIAL

**Supplemental File S1. WGBS sample names and experimental classifications**. List of the WGBS samples used in this study, including sample name, genotype, experimental lineage, generation, and experimental phase. Treatment labels in G5 reflect the preceding treatment lineage, as plants were grown under control conditions during the recovery generations.

**Supplemental File S2. Complete hypermethylated and hypomethylated DMR dataset.** Coordinate-level catalogue of hypermethylated and hypomethylated regions identified in G1, G3, and G5. Records are classified by genotype, treatment lineage, methylation context (CG, CHG, or CHH), generation, and methylation direction. Reads were aligned to the TAIR10 reference genome, whereas DMR direction was defined relative to the corresponding WT Control comparison baseline.

**Supplemental Table S3. AlphaBeta fitted estimates and pairwise statistical comparisons.** Fitted intercept, methylation-gain (α), and methylation-loss (β) estimates are provided separately for the CG, CHG, and CHH contexts. The workbook also contains pairwise comparisons between genotypes and treatments, including estimated differences, pooled standard errors, and nominal *P*-values for α and β. Nominal *P* < 0.05 is highlighted.

**Supplemental File S4. Summary of persistent and switching DMRs across G1, G3, and G5.** Numbers of DMR regions classified as persistent or switching, stratified by genotype, treatment, and methylation context. Persistent DMRs retained the same hypermethylated or hypomethylated direction in G1, G3, and G5. Switcher DMRs were detected across all three sampled generations, but changed methylation direction in at least one generation.

**Supplemental File S5. Literature-supported persistent and switching DMR candidates.** Literature-guided prioritisation of genes and transposable elements associated with persistent or switching DMRs. The persistent section contains the curated candidate-gene set, direct promoter or gene-body DMR associations, candidate-selection summaries, supporting references, and the candidate universe. The switcher section contains directly associated genes and transposable elements, recurrent genomic loci, gene–TE co-overlapping loci, prioritisation tiers, and supporting references.

**Supplemental File S6. Integrated DMR and RNA-seq data for candidate genes.** Matched WGBS and RNA-seq data underlying the persistent- and switcher-candidate heatmaps. Separate worksheets provide DMR methylation state, promoter or gene-body association, RNA-seq log_2_ fold change, DESeq2-adjusted *P*-value, and expression-significance classification for each matched genotype, treatment, and generation. Significance was defined as adjusted *P* < 0.05.

**Supplemental File S7. Persistent TE-associated DMRs and expression of nearby genes.** Condition-matched integration of persistent TE-associated DMRs with significant RNA-seq responses of nearby genes. The detailed worksheet reports TE identity, family and superfamily, genomic coordinates, nearby gene, TE–gene distance and positional relationship, genotype, treatment, generation, TE methylation state, gene-expression log fold change, and adjusted *P*-value. Additional worksheets summarise unique TEs, genes, TE–gene pairs, and matched events by TE superfamily.

**Supplemental File S8. Validation of current DMRs against previous Arabidopsis stress-methylome studies.** Coordinate-level comparison of the current G1–G3–G5 DMR trajectory atlas with a compiled atlas of previously reported Arabidopsis stress-associated DMRs. Overlaps were required to occur in the same methylation context. The workbook contains the current and published DMR atlases; individual current–published overlap relationships; summaries by genotype, treatment, context, trajectory class, and published study; enrichment tests with raw and Benjamini–Hochberg-adjusted *P*-values; and nonexclusive promoter, gene-body, TE, and intergenic annotations. Each row in the overlap table represents an overlap relationship rather than a unique DMR; therefore, one current DMR may be linked to multiple published DMRs or studies.

**Supplemental File S9. Independent validation against the Rudolf et al. (2021) WT–gsnor1-3 methylome dataset.** Coordinate- and direction-based comparison of current G1 *gsnor-ko* Control DMRs with published *gsnor1-3*-versus-WT DMRs from Rudolf et al. (2021). The workbook contains all pairwise coordinate overlaps; classifications of current DMRs according to overlap and directional concordance; the concordantly validated G1 anchor set; reciprocal summaries using either current or published DMRs as the denominator; and audits of one-to-many and many-to-many overlap relationships. Directional agreement was defined by concordance between the current and published hypermethylated or hypomethylated state. The published DMR table did not report individual CG, CHG, or CHH contexts; consequently, methylation-context agreement could not be assessed.

## DATA AVAILABILITY

The raw WGBS data supporting this study have been deposited in ArrayExpress under accession **E-MTAB-17204**. Processed data and associated supporting materials, including supplemental material and FastQC quality-control reports, have been deposited in Zenodo – https://doi.org/10.5281/zenodo.22960793.

## Abbreviations

AI, artificial intelligence; ANOVA, analysis of variance; CTAB, cetyltrimethylammonium bromide; DMR, differentially methylated region; G1–G5, generations 1–5; *gsnor-ko*, GSNOR-deficient *gsnor1-3* mutant; GSNOR, S-nitrosoglutathione reductase; LINE, long interspersed nuclear element; LTR, long terminal repeat; PCA, principal component analysis; PCR, polymerase chain reaction; RC, rolling circle; RdDM, RNA-directed DNA methylation; RNA-seq, RNA sequencing; SINE, short interspersed nuclear element; TAIR10, The Arabidopsis Information Resource 10 reference genome; TE, transposable element; WGBS, whole-genome bisulfite sequencing; WT, wild type.

## Notes

https://doi.org/10.5281/zenodo.22960793

### Competing Interest Statement

The authors have declared no competing interest.

