## Supplementary material for "GSNOR-dependent nitric oxide homeostasis shapes methylome persistence after multi-generational exposure to future climate conditions in *Arabidopsis thaliana*": Behl et al_Supplemental Figures

**Supplemental Table S1**

**Pairwise comparisons of AlphaBeta methylation-gain and methylation-loss parameters**

Pairwise comparisons of methylation-gain (α) and methylation-loss (β) parameters estimated using the AlphaBeta ABneutral model, shown separately for (A) CG, (B) CHG and (C) CHH methylation. Comparisons include treatments versus the corresponding genotype control, mutant versus WT controls, and genotype comparisons under matched treatments. Values are nominal P values; no multiple-testing adjustment is applied in this table. Bold red values indicate P < 0.05. Values below 0.001 are reported as <0.001. WT, wild-type Arabidopsis thaliana Col-0; *gsnor-ko*, the GSNOR-deficient *gsnor1-3* mutant. CO₂, elevated carbon dioxide; Ozone, elevated ozone; Combination, combined environmental treatment.

**A CG methylation**

| **Comparison** | **P (α)** | **P (β)** |
| --- | --- | --- |
| WT Drought vs WT Control | 0.433 | 0.392 |
| WT CO₂ vs WT Control | 0.066 | **0.049** |
| WT Ozone vs WT Control | 0.241 | 0.248 |
| WT Heat vs WT Control | 0.248 | 0.498 |
| WT Combination vs WT Control | **0.012** | 0.086 |
| *gsnor-ko* Control vs WT Control | **0.015** | 0.344 |
| *gsnor-ko* Drought vs *gsnor-ko* Control | 0.063 | 0.057 |
| *gsnor-ko* CO₂ vs *gsnor-ko* Control | **0.024** | 0.068 |
| *gsnor-ko* Ozone vs *gsnor-ko* Control | **0.033** | **0.020** |
| *gsnor-ko* Heat vs *gsnor-ko* Control | **0.003** | **0.004** |
| *gsnor-ko* Combination vs *gsnor-ko* Control | 0.248 | 0.192 |
| *gsnor-ko* Drought vs WT Drought | 0.422 | 0.305 |
| *gsnor-ko* CO₂ vs WT CO₂ | **<0.001** | **<0.001** |
| *gsnor-ko* Ozone vs WT Ozone | 0.144 | **0.019** |
| *gsnor-ko* Heat vs WT Heat | **0.001** | **0.009** |
| *gsnor-ko* Combination vs WT Combination | 0.181 | 0.091 |

**B CHG methylation**

| **Comparison** | **P (α)** | **P (β)** |
| --- | --- | --- |
| WT Drought vs WT Control | 0.076 | **0.048** |
| WT CO₂ vs WT Control | 0.217 | 0.185 |
| WT Ozone vs WT Control | 0.223 | 0.200 |
| WT Heat vs WT Control | 0.051 | **0.036** |
| WT Combination vs WT Control | **0.049** | 0.078 |
| *gsnor-ko* Control vs WT Control | 0.064 | **0.010** |
| *gsnor-ko* Drought vs *gsnor-ko* Control | **0.030** | **0.028** |
| *gsnor-ko* CO₂ vs *gsnor-ko* Control | 0.280 | 0.412 |
| *gsnor-ko* Ozone vs *gsnor-ko* Control | **0.034** | **0.027** |
| *gsnor-ko* Heat vs *gsnor-ko* Control | 0.250 | 0.281 |
| *gsnor-ko* Combination vs *gsnor-ko* Control | 0.157 | 0.136 |
| *gsnor-ko* Drought vs WT Drought | 0.264 | 0.217 |
| *gsnor-ko* CO₂ vs WT CO₂ | 0.495 | 0.187 |
| *gsnor-ko* Ozone vs WT Ozone | 0.061 | **0.034** |
| *gsnor-ko* Heat vs WT Heat | 0.145 | 0.240 |
| *gsnor-ko* Combination vs WT Combination | **0.002** | **<0.001** |

**C CHH methylation**

| **Comparison** | **P (α)** | **P (β)** |
| --- | --- | --- |
| WT Drought vs WT Control | 0.256 | 0.315 |
| WT CO₂ vs WT Control | 0.261 | 0.347 |
| WT Ozone vs WT Control | 0.094 | 0.118 |
| WT Heat vs WT Control | 0.415 | 0.420 |
| WT Combination vs WT Control | 0.397 | 0.486 |
| *gsnor-ko* Control vs WT Control | 0.202 | 0.239 |
| *gsnor-ko* Drought vs *gsnor-ko* Control | 0.145 | 0.127 |
| *gsnor-ko* CO₂ vs *gsnor-ko* Control | 0.168 | 0.216 |
| *gsnor-ko* Ozone vs *gsnor-ko* Control | 0.258 | 0.288 |
| *gsnor-ko* Heat vs *gsnor-ko* Control | 0.374 | 0.397 |
| *gsnor-ko* Combination vs *gsnor-ko* Control | 0.102 | 0.120 |
| *gsnor-ko* Drought vs WT Drought | 0.372 | 0.314 |
| *gsnor-ko* CO₂ vs WT CO₂ | 0.452 | 0.437 |
| *gsnor-ko* Ozone vs WT Ozone | 0.136 | 0.157 |
| *gsnor-ko* Heat vs WT Heat | 0.434 | 0.482 |
| *gsnor-ko* Combination vs WT Combination | 0.219 | 0.189 |

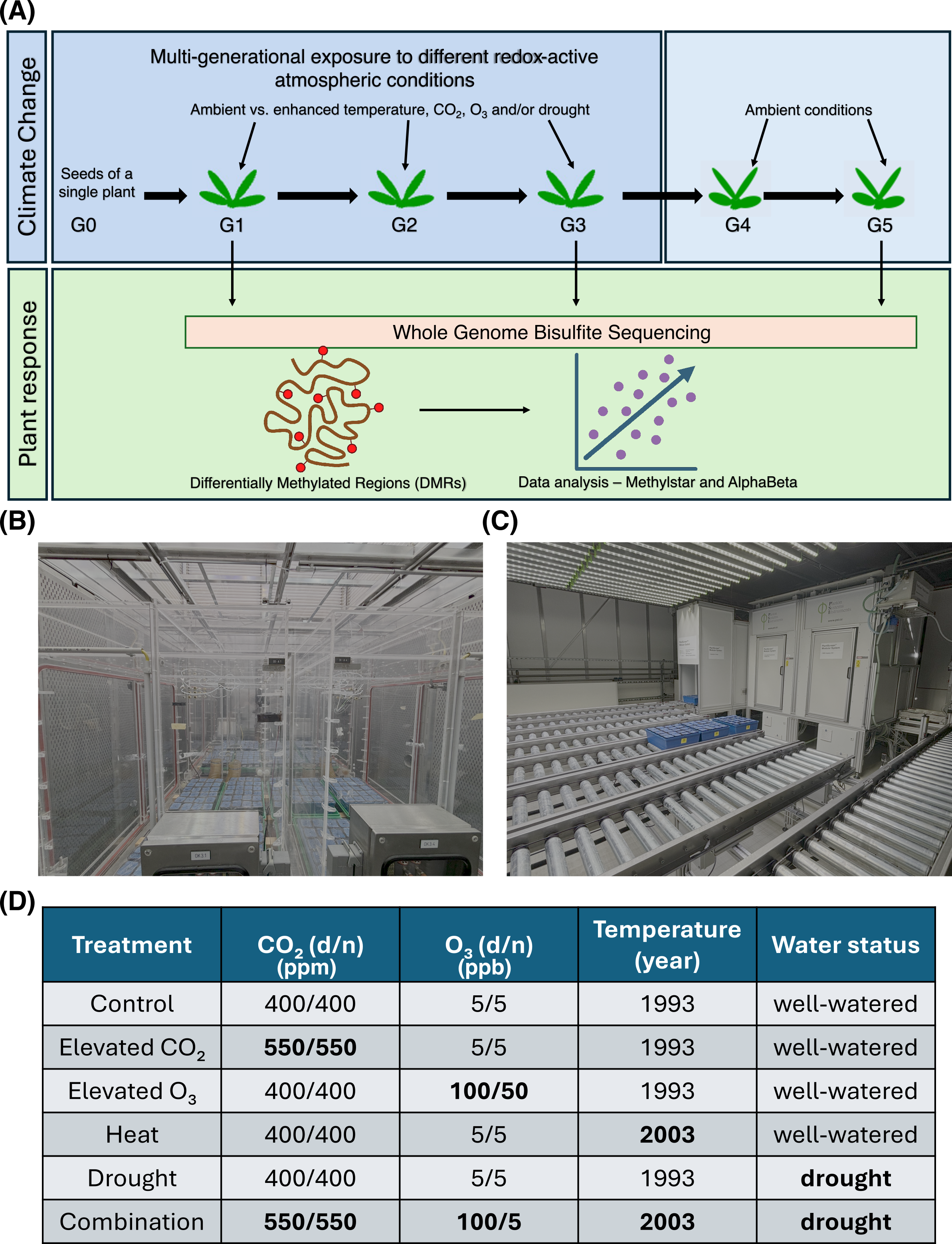

**Supplemental Figure S1 - Experimental design and facilities used for the multigenerational climate-stress experiment.** **(A)** Overview of the five-generation experiment in wild type (WT) and *gsnor-ko* *Arabidopsis thaliana*. Plants were propagated from a single parental plant within each genotype and exposed to control or altered temperature, CO₂, O₃ and/or water availability during G1–G3. G4 and G5 were grown under control conditions to assess recovery. Leaf samples from G1, G3 and G5 were analysed by whole-genome bisulfite sequencing, followed by methylation processing, DMR identification and AlphaBeta analysis. **(B)** Climate-simulation chambers used for plant cultivation and environmental treatments. **(C)** Automated plant-phenotyping facility used to assess plant responses. **(D)** Overview of the six environmental scenarios. Atmospheric concentrations are given separately for day and night (d/n). Temperature regimes are identified by the reference years 1993 and 2003.

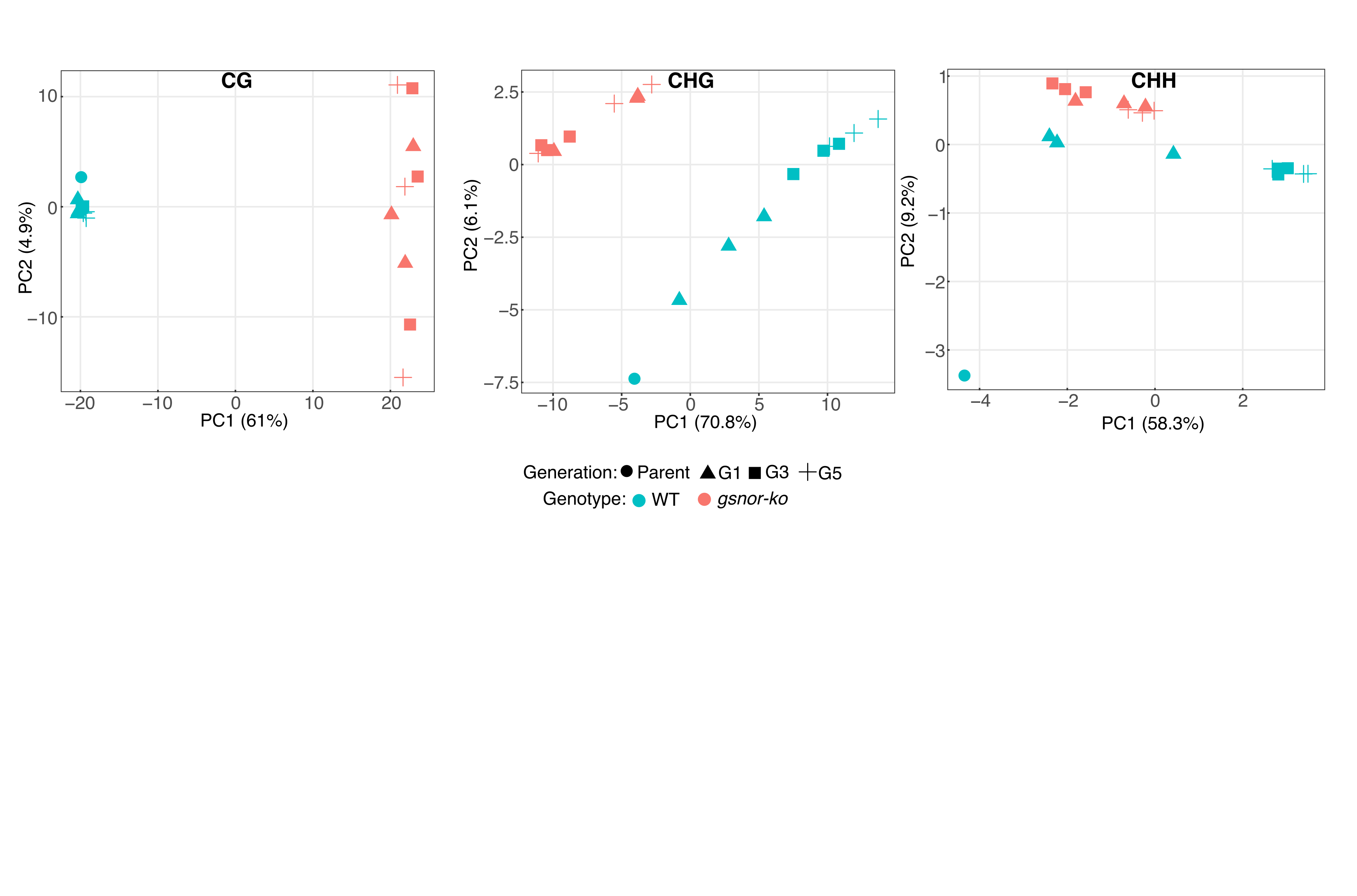

**Supplemental Figure S2 - Principal component analysis of control methylomes across generations.** Principal component analysis (PCA) of WT and *gsnor-ko* methylation profiles under control conditions, performed separately for CG, CHG and CHH sequence contexts. Each point represents an individual WGBS sample. Colours distinguish WT (turquoise) and *gsnor-ko* (salmon), while symbols indicate the parental sample, G1, G3 and G5. PC1 and PC2 explain 61.0% and 4.9% of the variation in CG, 70.8% and 6.1% in CHG, and 58.3% and 9.2% in CHH, respectively.

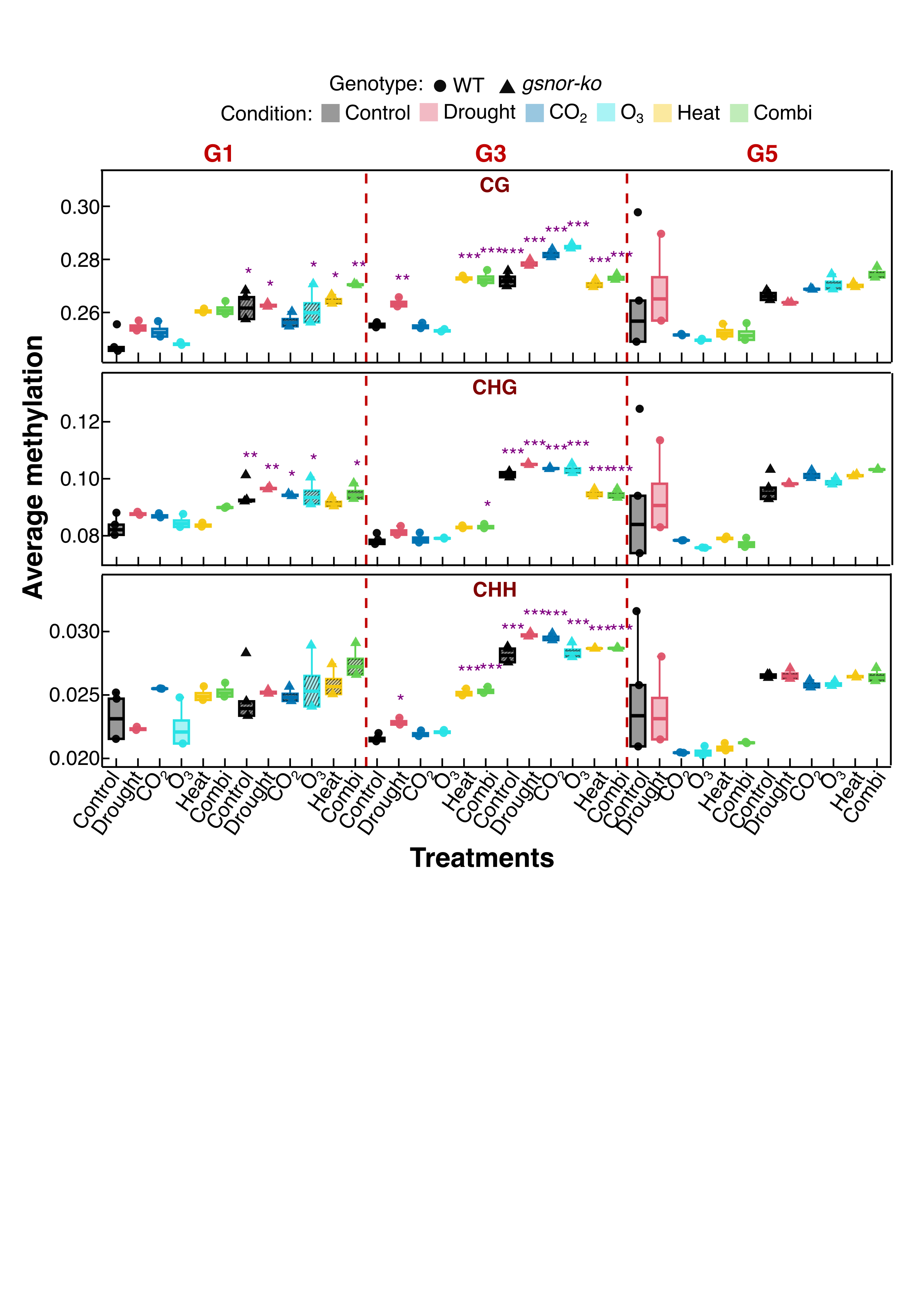

**Supplemental Figure S3 - Genome-wide average methylation across genotypes, treatments and generations.** Average methylation in the CG, CHG and CHH sequence contexts in WT and *gsnor-ko* plants sampled at G1, G3 and G5. Colours distinguish treatments; circles represent WT biological replicates and triangles represent *gsnor-ko* replicates. WT boxes are plain, and *gsnor-ko* boxes are striped. Boxes show the median and interquartile range, with whiskers extending to the most extreme observations within 1.5 times the interquartile range. Three biological replicates were analysed per control group and two per treatment group. Vertical dashed lines separate generations. Genotype, treatment, and their interaction were assessed using two-factor ANOVA, separately for each methylation context and generation. Asterisks indicate multiplicity-adjusted comparisons against WT Control within the corresponding generation: P < 0.05 (*), P < 0.01 (**) and P < 0.001 (***). G5 treatment labels indicate ancestral treatment lineages following two generations under control conditions.

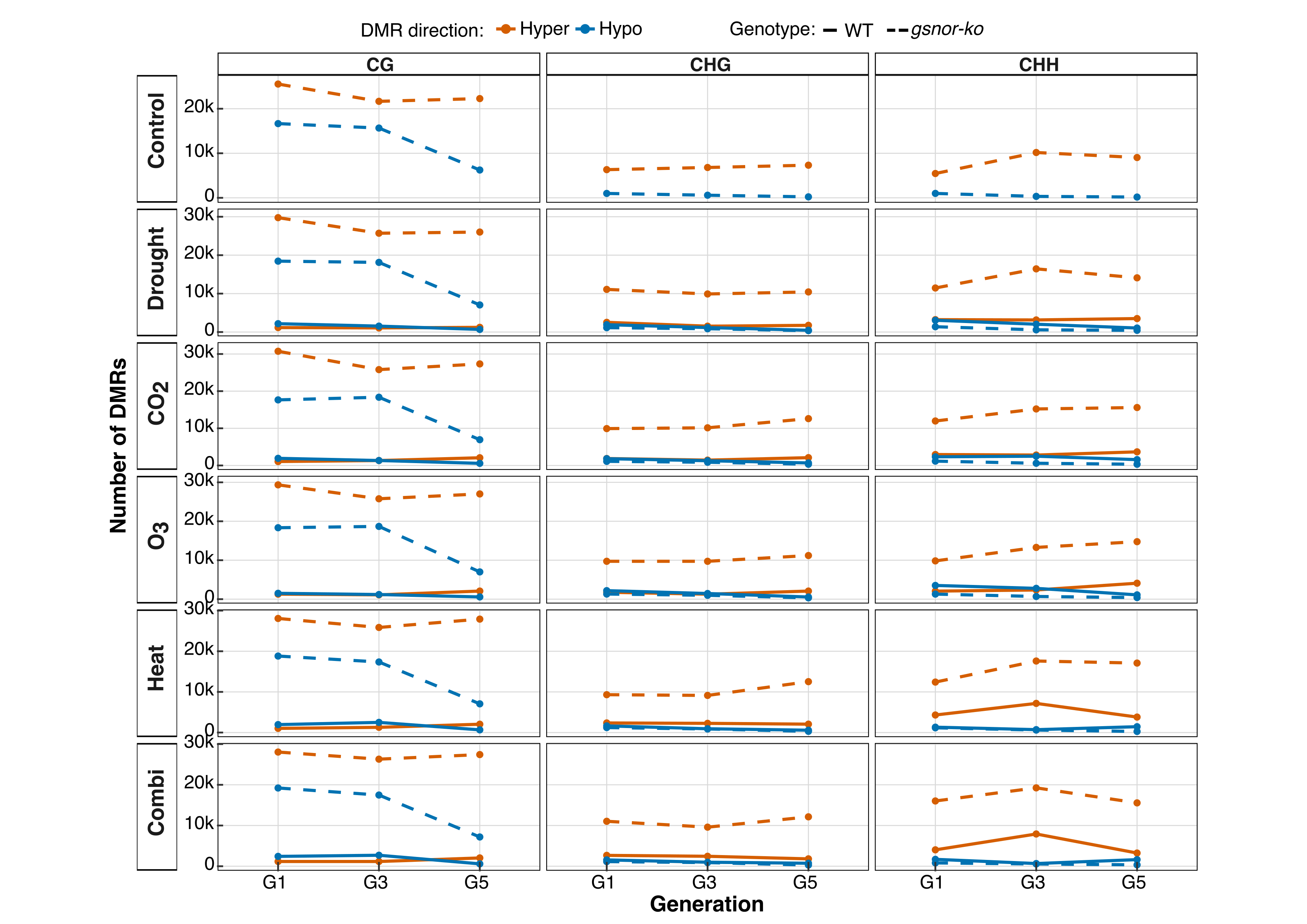

**Supplemental Figure S4 - Generational trajectories of hypermethylated and hypomethylated DMR counts.** Numbers of hypermethylated (orange) and hypomethylated (blue) DMRs across G1, G3 and G5, separated by treatment lineage and methylation context. Solid lines represent WT and dashed lines represent *gsnor-ko*. All differential methylation states were defined relative to WT Control within the corresponding generation. The Control row therefore shows the baseline *gsnor-ko*-versus-WT comparison. DMRs were identified using jDMRgrid with 200-bp windows, a 50-bp step, a minimum of 10 cytosines per context and replicate, and consensus = 1. Points show DMR counts, and connecting lines illustrate changes across sampling points. G5 labels indicate ancestral treatment lineages following two generations under control conditions.

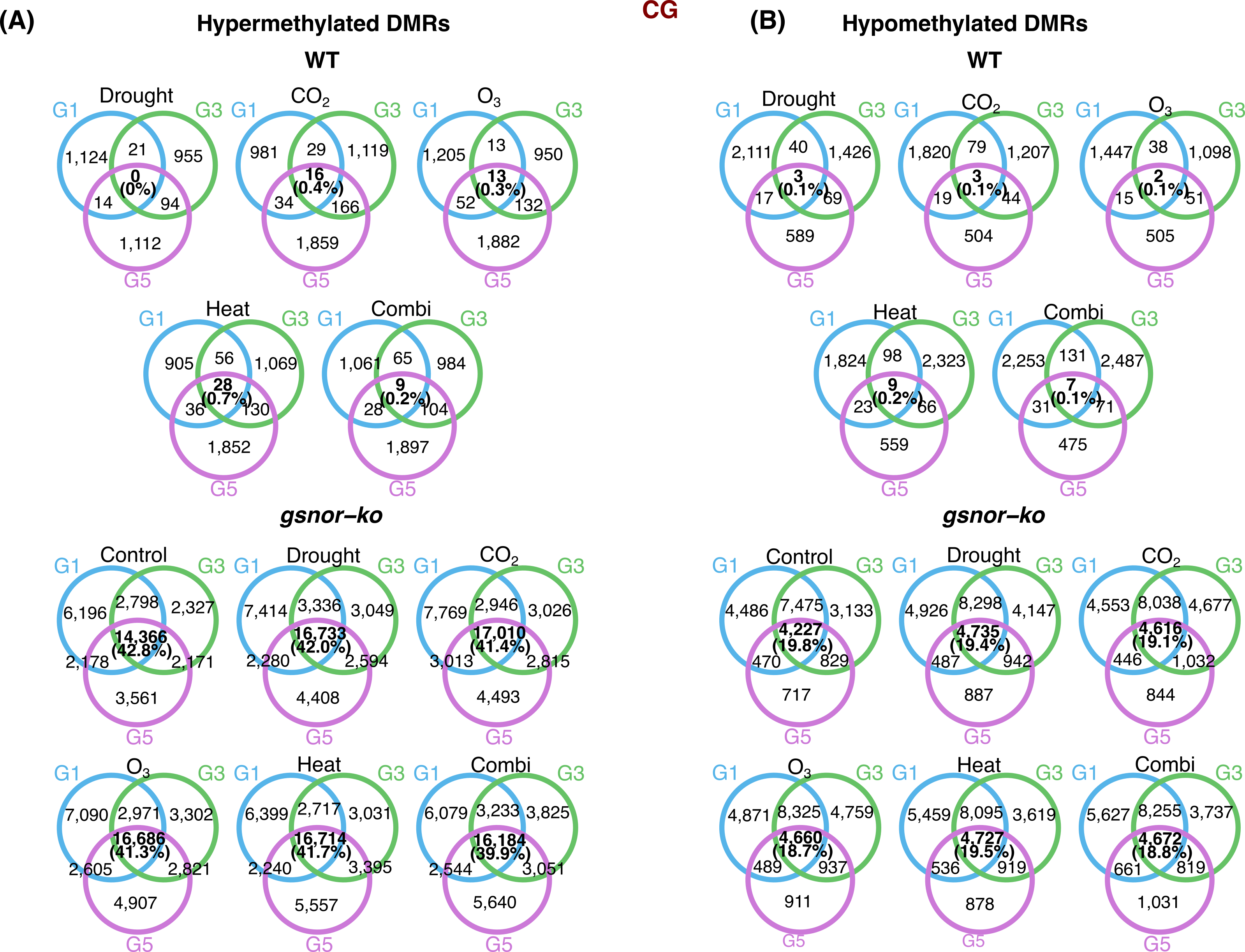

**Supplemental Figure S5 - Overlap of CG DMRs across G1, G3 and G5.** Three-set Venn diagrams showing overlaps among CG DMRs detected at G1, G3 and G5 within each genotype and treatment lineage. Hypermethylated **(A)** and hypomethylated **(B)** DMRs are shown separately. Blue, green and purple outlines represent G1, G3 and G5, respectively. All DMRs were defined relative to WT Control within the corresponding generation; the *gsnor-ko* Control panels represent baseline genotype-associated differences. Overlapping genomic intervals were harmonised into consensus regions using a minimum overlap of 1 bp before calculating set intersections. Numbers indicate regions unique to or shared among the indicated generations. Bold central labels show the number detected in all three generations in the same methylation direction and its percentage of the three-generation union. Percentages are rounded. G5 labels refer to ancestral treatments followed by two generations under control conditions.

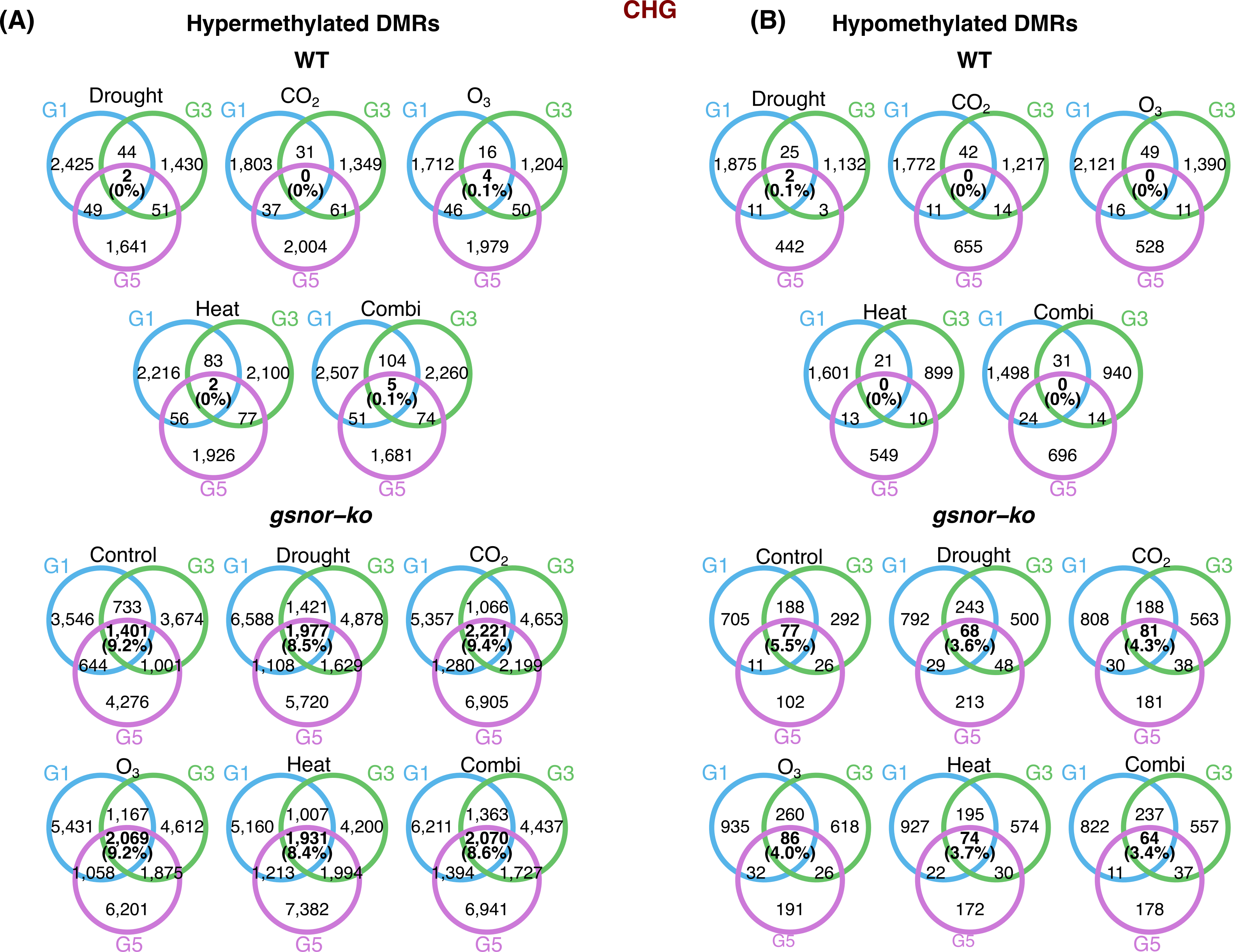

**Supplemental Figure S6 - Overlap of CHG DMRs across G1, G3 and G5.** Three-set Venn diagrams showing overlaps among CHG DMRs detected at G1, G3 and G5 within each genotype and treatment lineage, separately for hypermethylated **(A)** and hypomethylated **(B)** regions. Blue, green and purple outlines represent G1, G3 and G5, respectively. DMR direction was defined relative to WT Control within the corresponding generation. Overlapping intervals were harmonised into consensus regions using a minimum overlap of 1 bp. Numbers indicate regions unique to or shared among generations. Bold central labels show the number detected in all three generations in the same methylation direction and its percentage of the three-generation union. Percentages are rounded. The *gsnor-ko* Control panels show baseline genotype-associated differences, and G5 treatment labels indicate ancestral treatment lineages following two generations under control conditions.

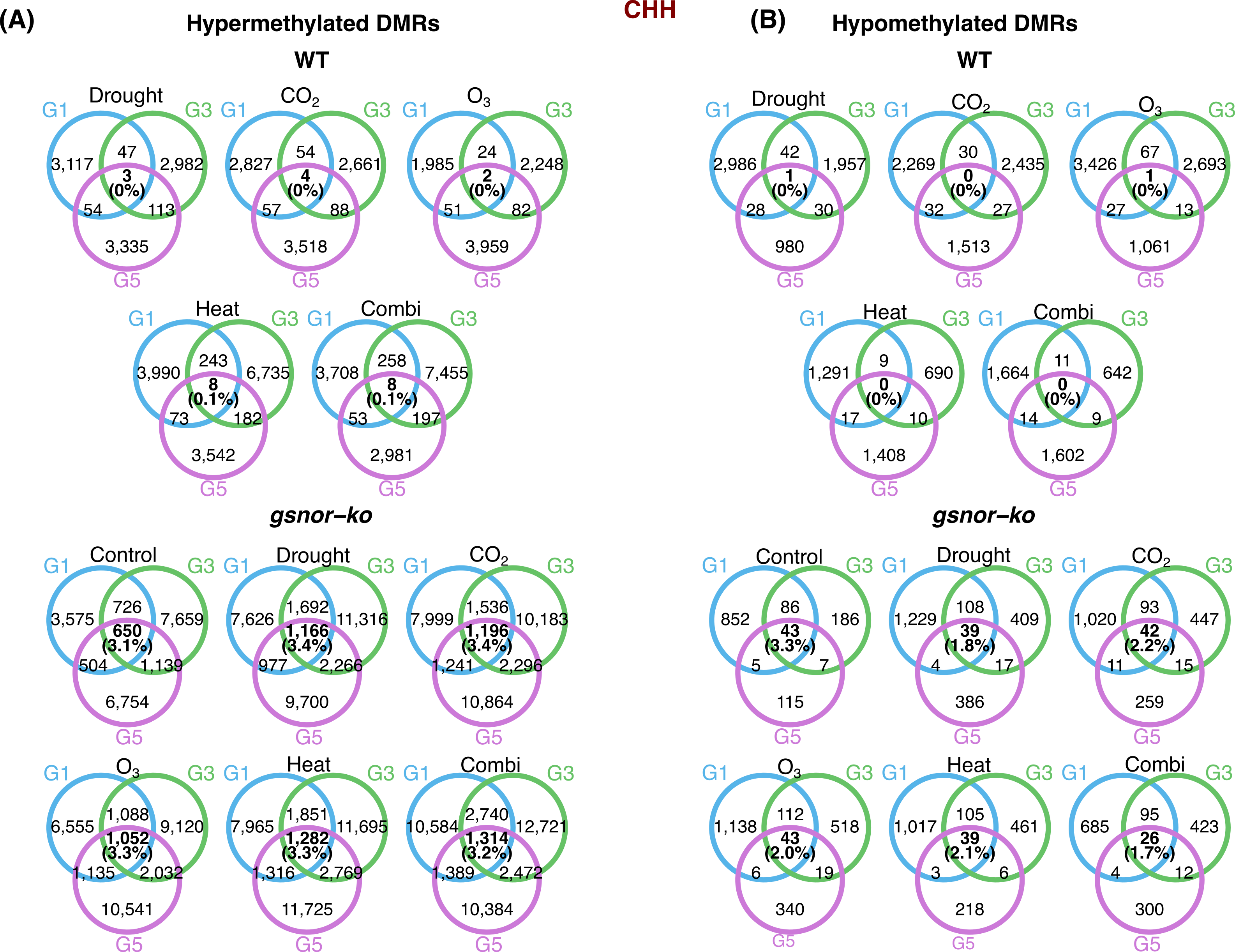

**Supplemental Figure S7 - Overlap of CHH DMRs across G1, G3 and G5.** Three-set Venn diagrams showing overlaps among CHH DMRs detected at G1, G3 and G5 within each genotype and treatment lineage, separately for hypermethylated **(A)** and hypomethylated **(B)** regions. Blue, green and purple outlines represent G1, G3 and G5, respectively. DMR direction was defined relative to WT Control within the corresponding generation. Overlapping intervals were harmonised into consensus regions using a minimum overlap of 1 bp. Numbers indicate regions unique to or shared among generations. Bold central labels show the number detected in all three generations in the same methylation direction and its percentage of the three-generation union. Percentages are rounded. The *gsnor-ko* Control panels show baseline genotype-associated differences, and G5 treatment labels indicate ancestral treatment lineages following two generations under control conditions.

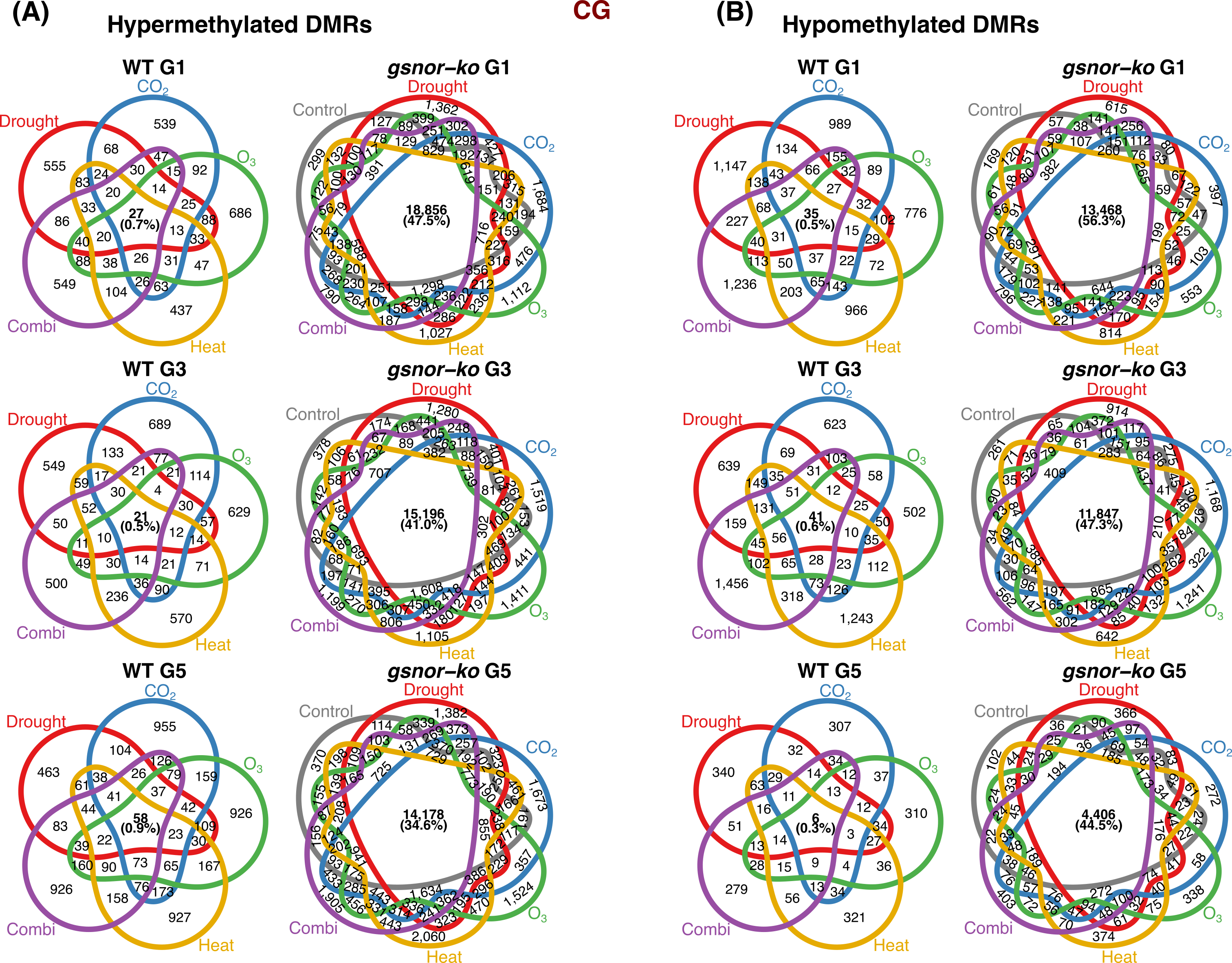

**Supplemental Figure S8 - Overlap of CG DMRs among treatment sets within each generation.** Venn diagrams showing overlaps among CG DMR sets in WT and *gsnor-ko* at G1, G3 and G5, separately for hypermethylated **(A)** and hypomethylated **(B)** regions. WT diagrams include drought, elevated CO₂, elevated O₃, heat and combination treatments. The gsnor-ko diagrams additionally include the mutant Control comparison. All DMR sets were defined relative to WT Control within the corresponding generation. Overlapping intervals were harmonised into consensus regions using a minimum overlap of 1 bp. Numbers indicate regions unique to or shared among the displayed sets. Bold central labels show the number shared by all five WT sets or all six *gsnor-ko* sets, along with its percentage of the respective union. The mutant central intersection therefore includes regions that are also detected in the *gsnor-ko* Control. Percentages are rounded. G5 labels indicate ancestral treatment lineages following two generations under control conditions.

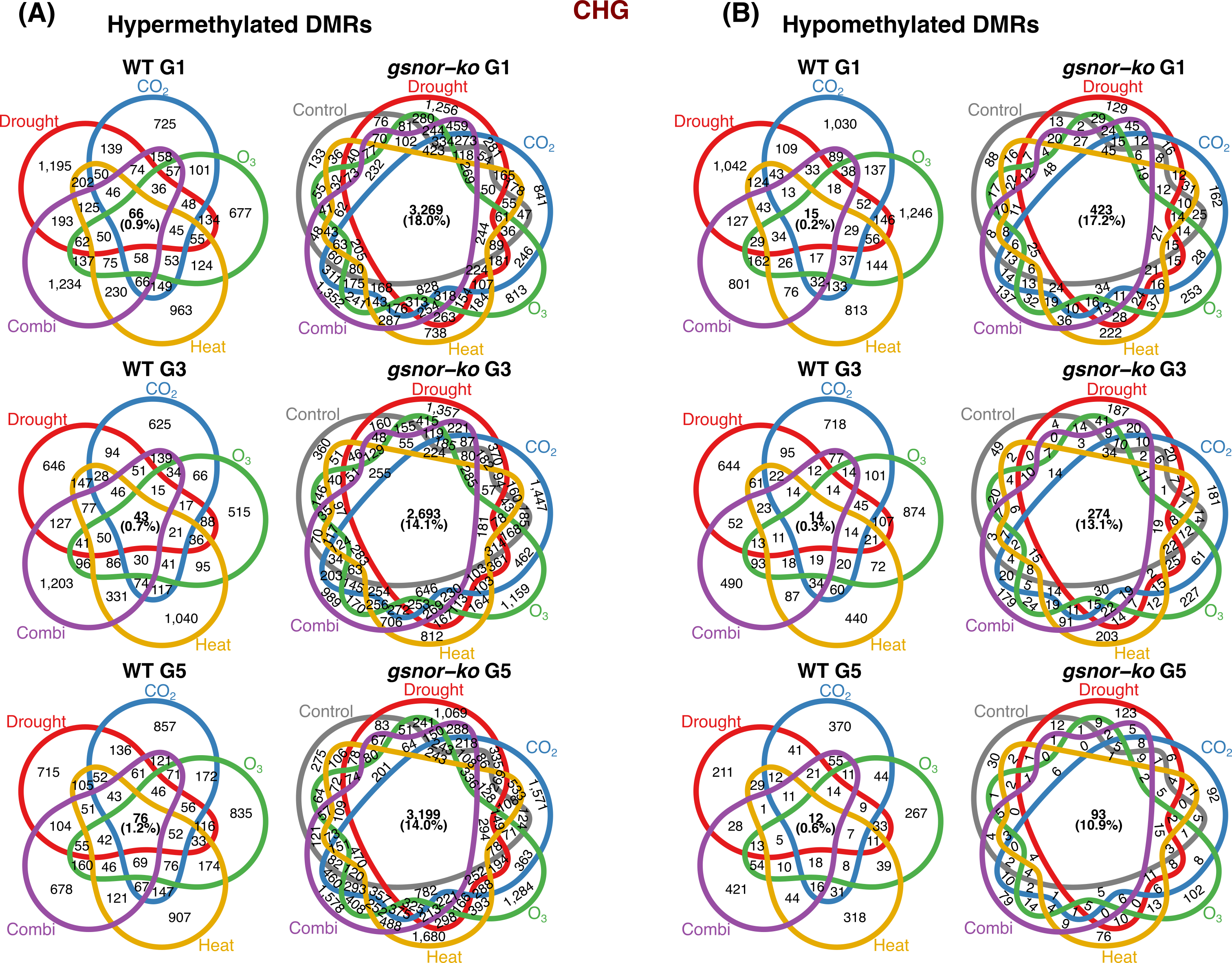

**Supplemental Figure S9 - Overlap of CHG DMRs among treatment sets within each generation.** Venn diagrams showing overlaps among CHG DMR sets in WT and *gsnor-ko* at G1, G3 and G5, separately for hypermethylated **(A)** and hypomethylated **(B)** regions. WT diagrams include the five environmental-treatment comparisons; *gsnor-ko* diagrams additionally include the mutant Control comparison. All comparisons used WT Control from the corresponding generation as the reference. Overlapping intervals were harmonised into consensus regions using a minimum overlap of 1 bp. Numbers indicate regions unique to or shared among the displayed sets. Bold central labels show the number shared by all five WT sets or all six *gsnor-ko* sets, along with its percentage of the respective union. The mutant central intersection therefore includes regions that are also detected in the *gsnor-ko* Control. Percentages are rounded. G5 labels indicate ancestral treatment lineages following two generations under control conditions.

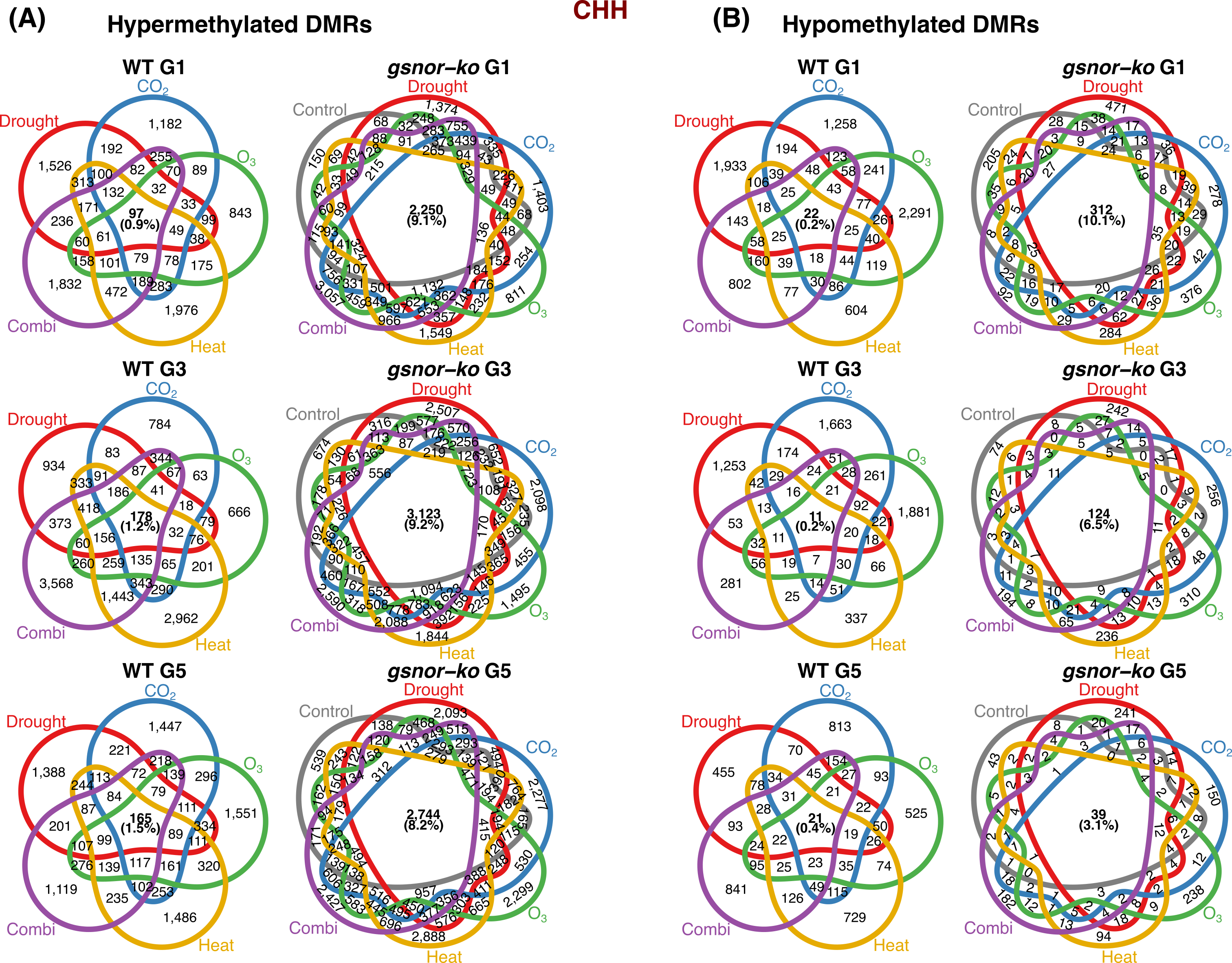

**Supplemental Figure S10 - Overlap of CHH DMRs among treatment sets within each generation.** Venn diagrams showing overlaps among CHH DMR sets in WT and *gsnor-ko* at G1, G3 and G5, separately for hypermethylated **(A)** and hypomethylated **(B)** regions. WT diagrams include the five environmental-treatment comparisons; *gsnor-ko* diagrams additionally include the mutant Control comparison. All comparisons used WT Control from the corresponding generation as the reference. Overlapping intervals were harmonised into consensus regions using a minimum overlap of 1 bp. Numbers indicate regions unique to or shared among the displayed sets. Bold central labels show the number shared by all five WT sets or all six *gsnor-ko* sets, along with its percentage of the respective union. The mutant central intersection therefore includes regions that are also detected in the *gsnor-ko* Control. Percentages are rounded. G5 labels indicate ancestral treatment lineages following two generations under control conditions.

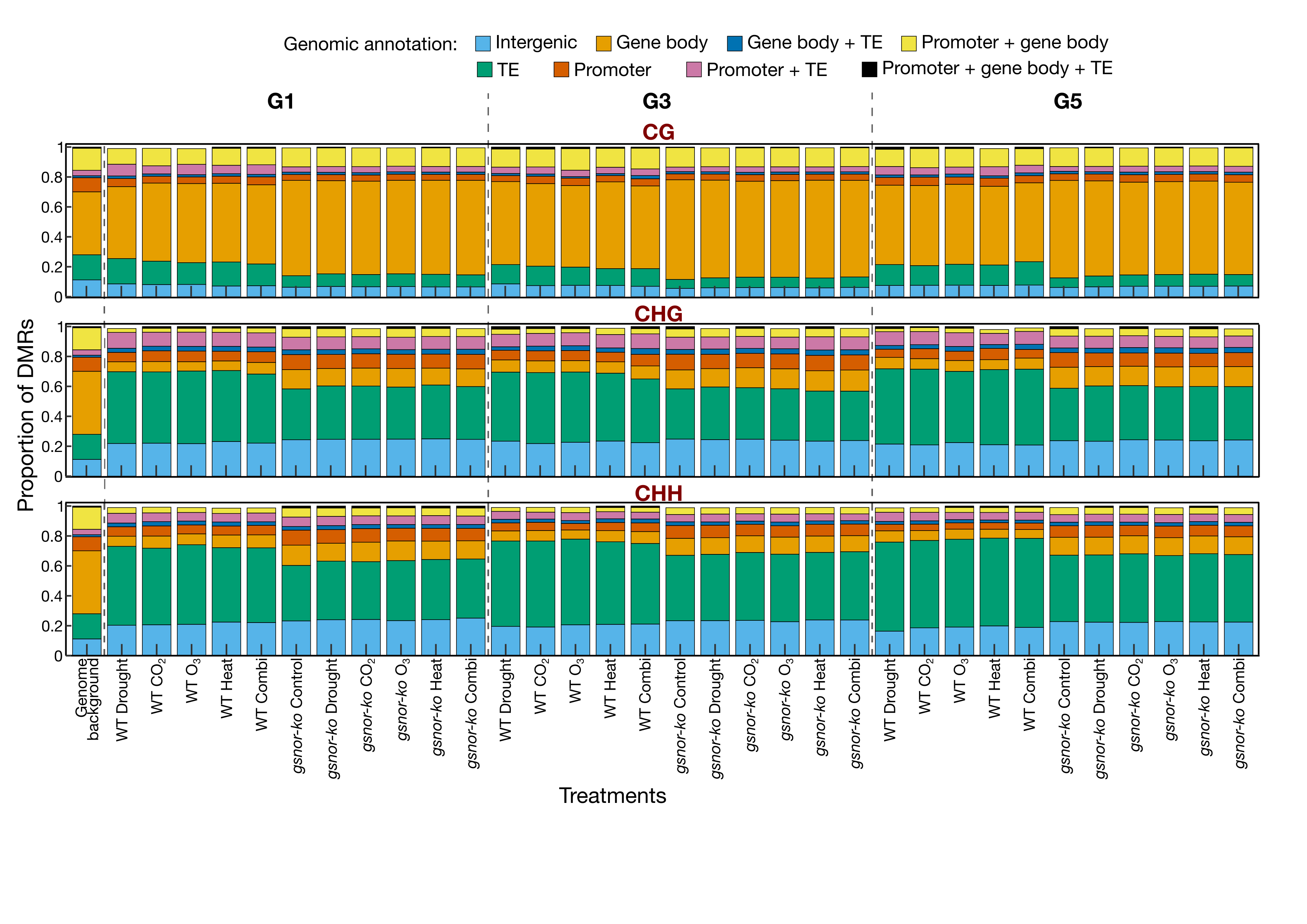

**Supplemental Figure S11 - Genomic distribution of differentially methylated regions across generations.** Proportions of DMRs overlapping genomic features in WT and *gsnor-ko* plants across treatments and generations, shown separately for CG, CHG and CHH sequence contexts. Hypermethylated and hypomethylated regions are combined. Regions were annotated against TAIR10 gene bodies, transposable elements (TEs) and promoters, defined as 1 kb upstream of the transcription start site. Each region was assigned to one of eight mutually exclusive annotation classes representing intergenic regions or the indicated combinations of feature overlaps. The genome-background bar shows annotation proportions for genome-wide 200-bp windows generated at 50-bp intervals across the five nuclear chromosomes. Other bars show proportions within each genotype–treatment–generation DMR set. Differential methylation was defined relative to WT Control within the corresponding generation. Dashed lines separate generations. G5 treatment labels indicate ancestral treatment lineages following two generations under control conditions.

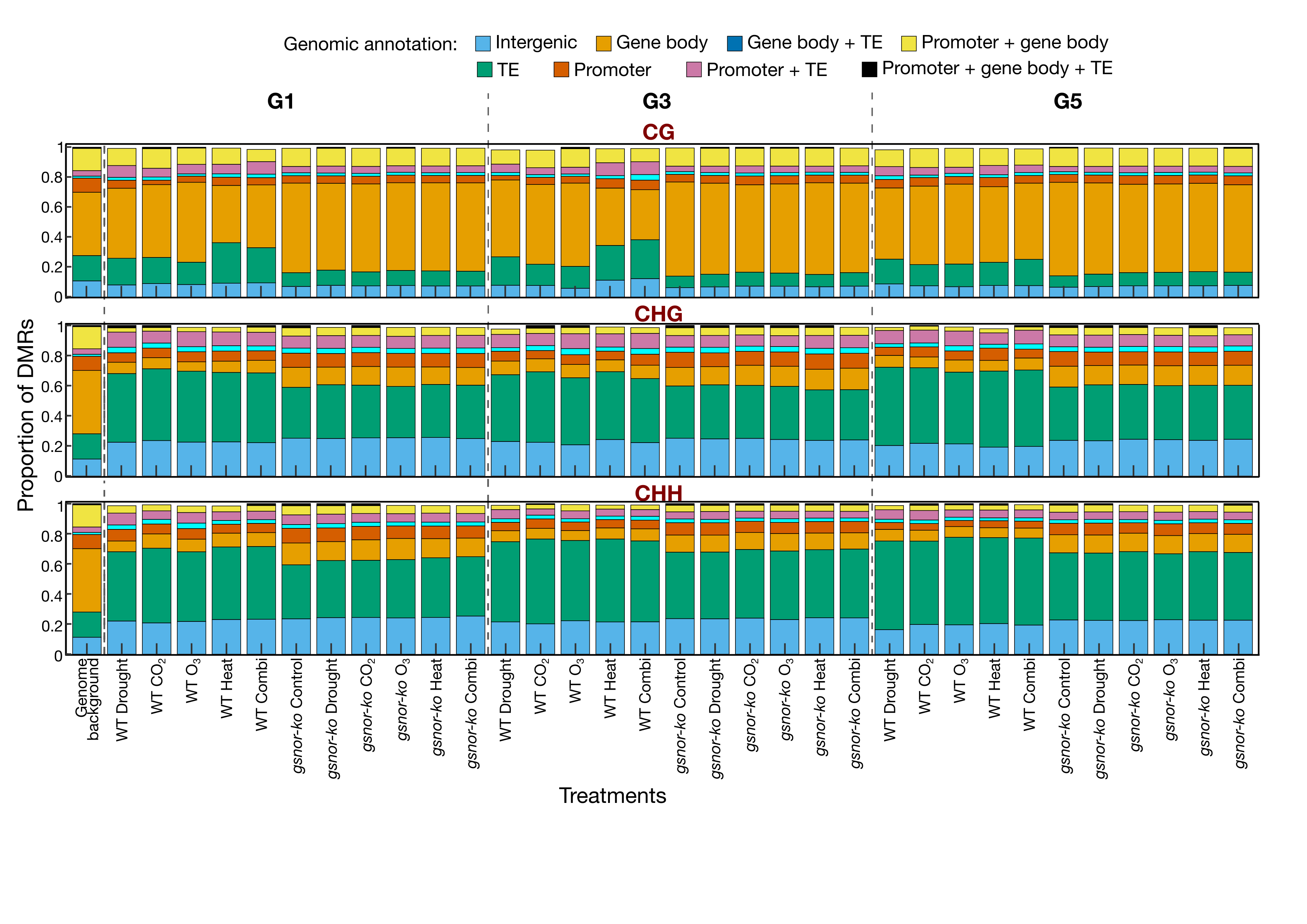

**Supplemental Figure S12 - Genomic distribution of hypermethylated DMRs across generations.** Proportions of hypermethylated DMRs assigned to genomic annotation classes in WT and *gsnor-ko* plants across G1, G3 and G5, shown separately for CG, CHG and CHH contexts. Hypermethylation was defined relative to WT Control within the corresponding generation. TAIR10 gene-body, TE and promoter overlaps were combined into eight mutually exclusive classes; promoters were defined as 1 kb upstream of the transcription start site. Intergenic regions overlapped none of these features. The genome-background bar represents annotation proportions for genome-wide 200-bp windows generated at 50-bp intervals across the five nuclear chromosomes. DMR proportions were calculated separately within each displayed comparison. Dashed lines separate generations. G5 labels indicate ancestral treatment lineages following two generations under control conditions.

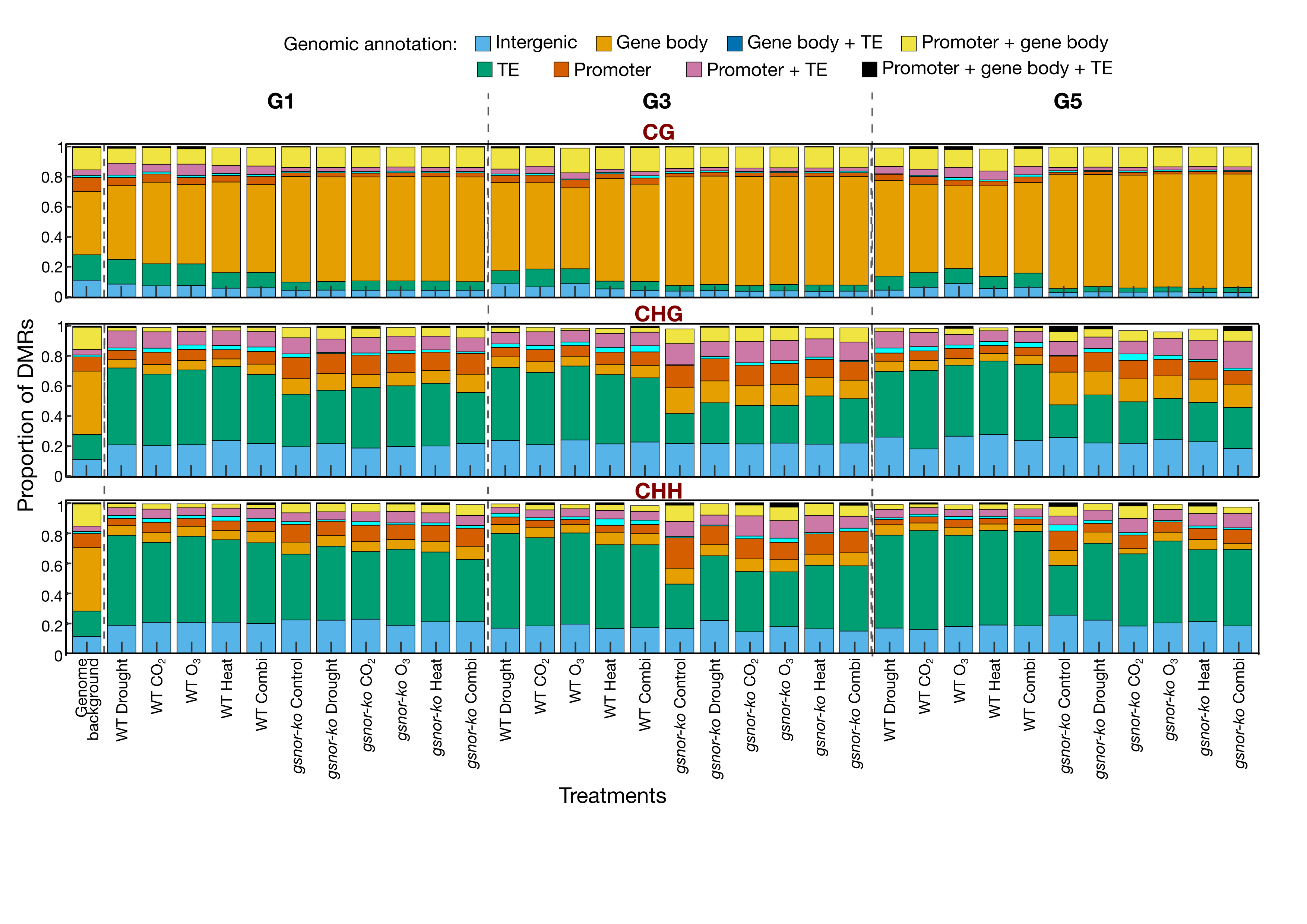

**Supplemental Figure S13 - Genomic distribution of hypomethylated DMRs across generations.** Proportions of hypomethylated DMRs assigned to genomic annotation classes in WT and *gsnor-ko* plants across G1, G3 and G5, shown separately for CG, CHG and CHH contexts. Hypomethylation was defined relative to WT Control within the corresponding generation. TAIR10 gene-body, TE and promoter overlaps were combined into eight mutually exclusive classes; promoters were defined as 1 kb upstream of the transcription start site. Intergenic regions overlapped none of these features. The genome-background bar represents annotation proportions for genome-wide 200-bp windows generated at 50-bp intervals across the five nuclear chromosomes. DMR proportions were calculated separately within each displayed comparison. Dashed lines separate generations. G5 labels indicate ancestral treatment lineages following two generations under control conditions.

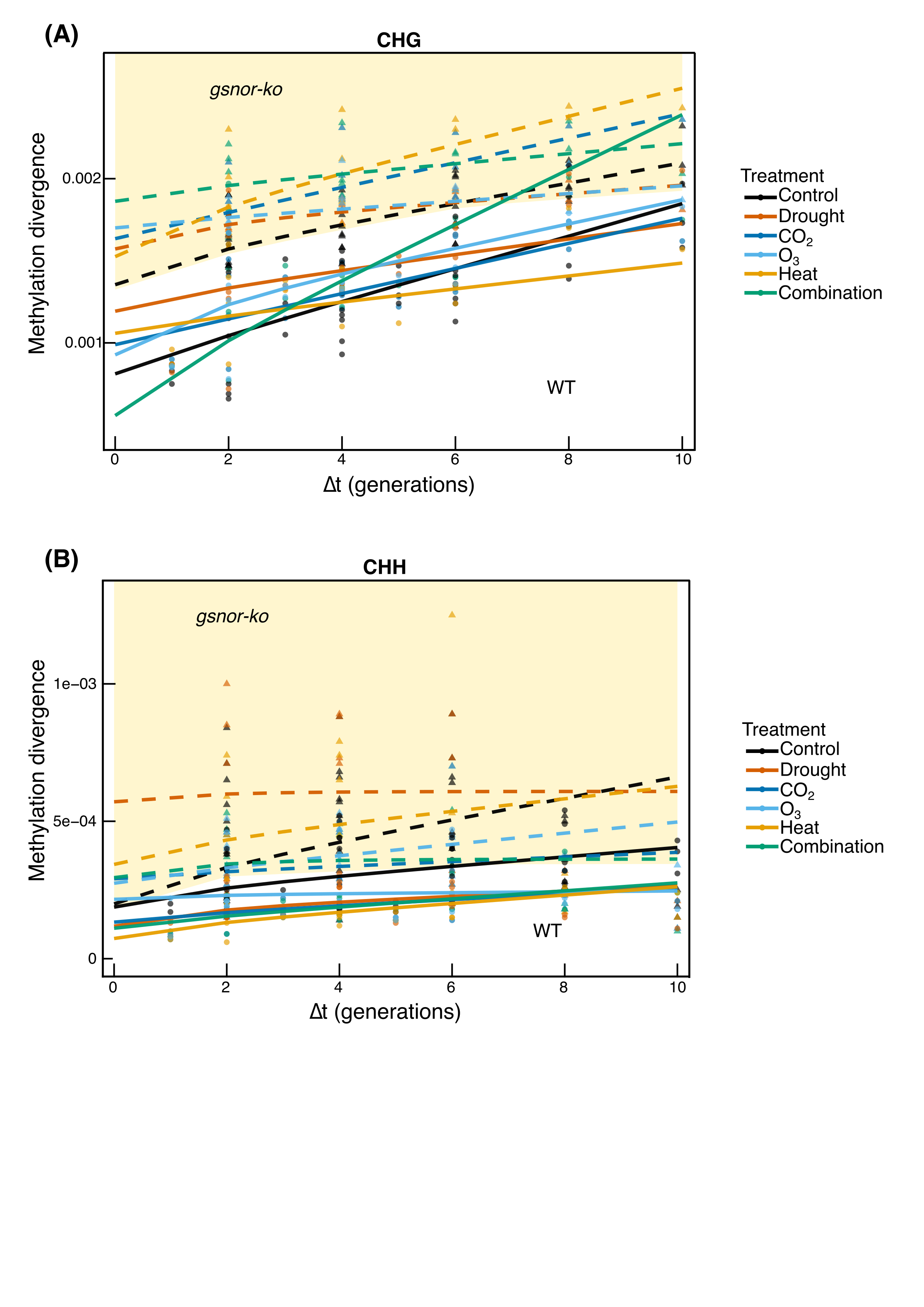

**Supplemental Figure S14 - AlphaBeta analysis of non-CG methylation divergence.** Observed methylation divergence and AlphaBeta-fitted trajectories in **(A)** CHG and **(B)** CHH sequence contexts across WT and *gsnor-ko* treatment lineages. Circles and solid lines represent WT observations and fitted trajectories, respectively; triangles and dashed lines represent *gsnor-ko*. Colours distinguish treatments. Δt denotes the summed generational distance separating two sampled individuals through their common ancestor. Methylation-gain (α) and methylation-loss (β) parameters were estimated using the ABneutral model with the experimental pedigree, a posteriorMaxFilter setting of 0.99 and 50 optimisation starts. Parameter uncertainty was assessed using 100 bootstrap iterations. Pale-yellow shading provides a visual highlight above a boundary placed just below the lowest fitted *gsnor-ko* trajectory; it does not represent a confidence interval or significance threshold.

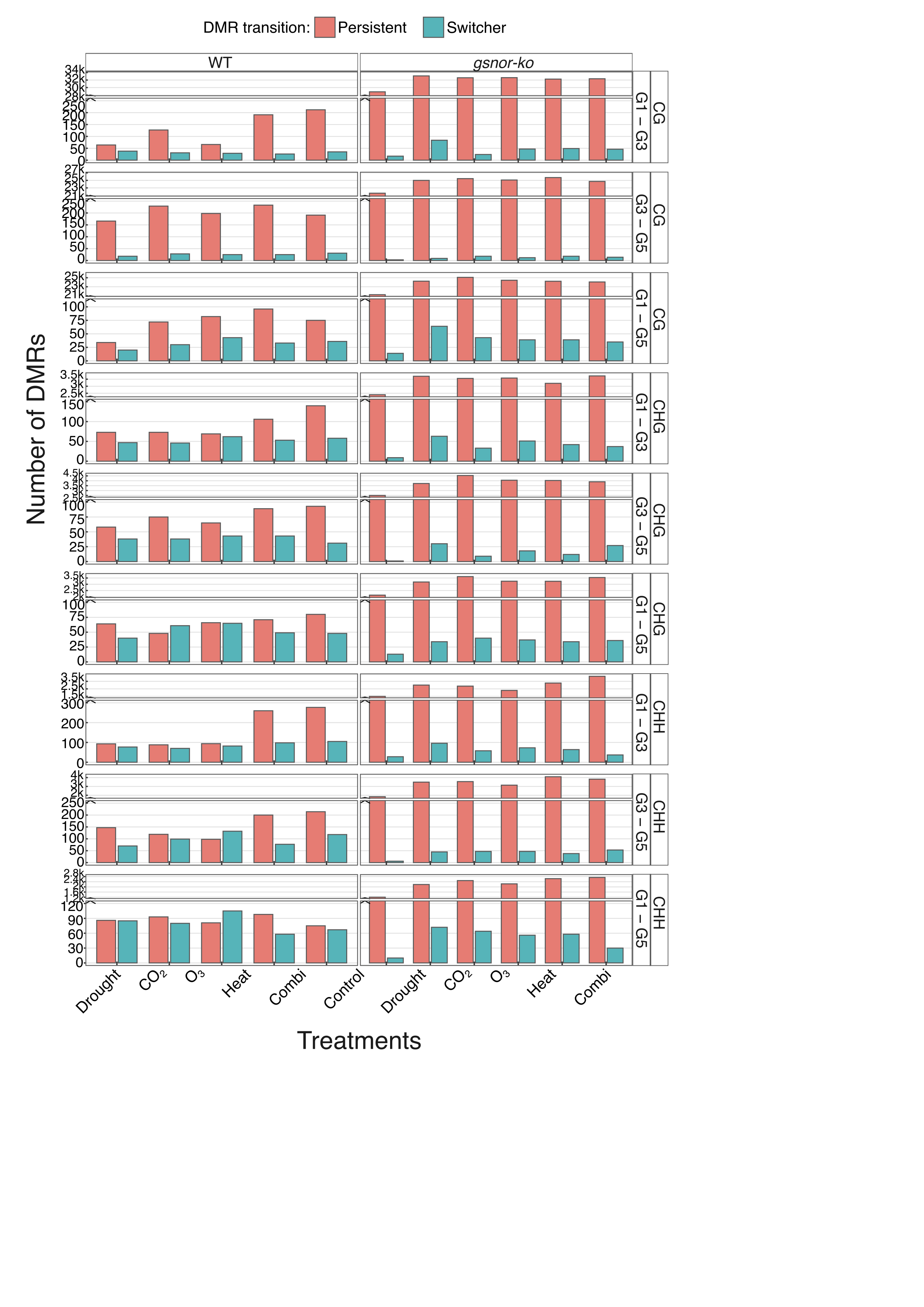

**Supplemental Figure S15 - Persistent and switching DMRs between pairs of sampled generations.** Numbers of DMRs retaining or reversing their differential methylation direction between G1–G3, G3–G5 and G1–G5, separated by genotype, treatment lineage and methylation context. Salmon bars indicate pairwise persistent regions, defined as hypermethylated or hypomethylated in the same direction at both sampling points. Turquoise bars indicate switchers, defined as hypermethylated at one sampling point and hypomethylated at the other. Classification considers only the two generations indicated and does not require the same behaviour in the third sampled generation. Differential methylation states were defined relative to WT Control within each generation. The *gsnor-ko* Control comparison represents baseline genotype-associated differences. Vertical-axis breaks accommodate differences in abundance. Bars show region counts rather than replicate means.

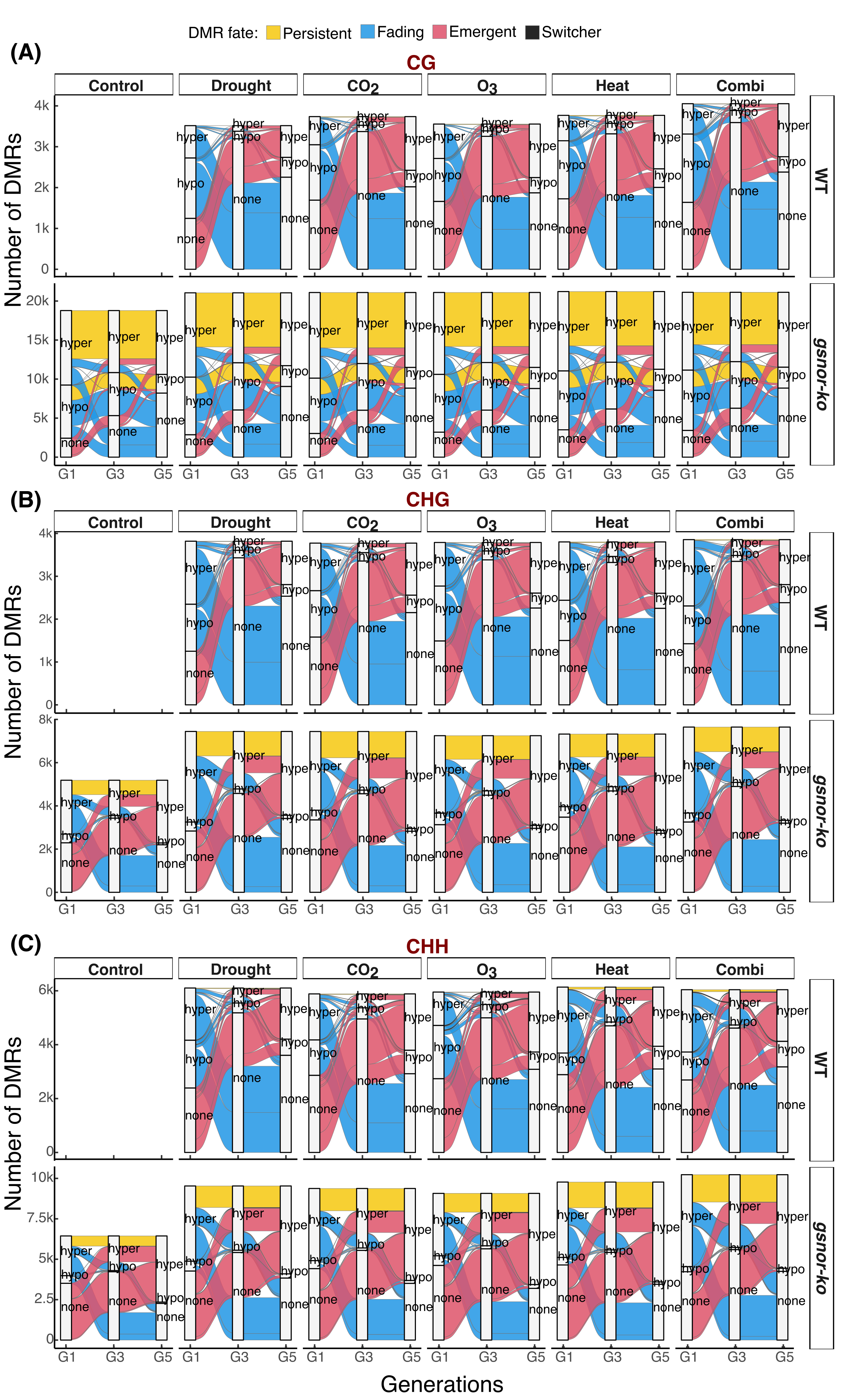

**Supplemental Figure S16 - Methylation-state trajectories across treatment and recovery generations.** Alluvial plots showing transitions among hypermethylated, hypomethylated and undetected differential methylation states across G1, G3 and G5 in **(A)** CG, **(B)** CHG and **(C)** CHH contexts. Panels are separated by genotype and treatment lineage. Ribbon widths represent the number of consensus genomic regions. Persistent regions (yellow) retained the same direction at all three sampling points; switchers (black) were detected at all three sampling points but changed direction at least once. Fading regions (blue) were detected in G1 but not G5, whereas emergent regions (pink) were absent in G1 and detected in G5. Ambiguous and other trajectories outside these four classes were excluded from the displayed flows. “None” indicates that no DMR was detected relative to WT Control in that generation, rather than an absence of DNA methylation. Vertical scales differ between genotypes. The WT Control panel is blank because WT Control served as the reference. G5 treatment labels indicate ancestral lineages following two generations under control conditions.

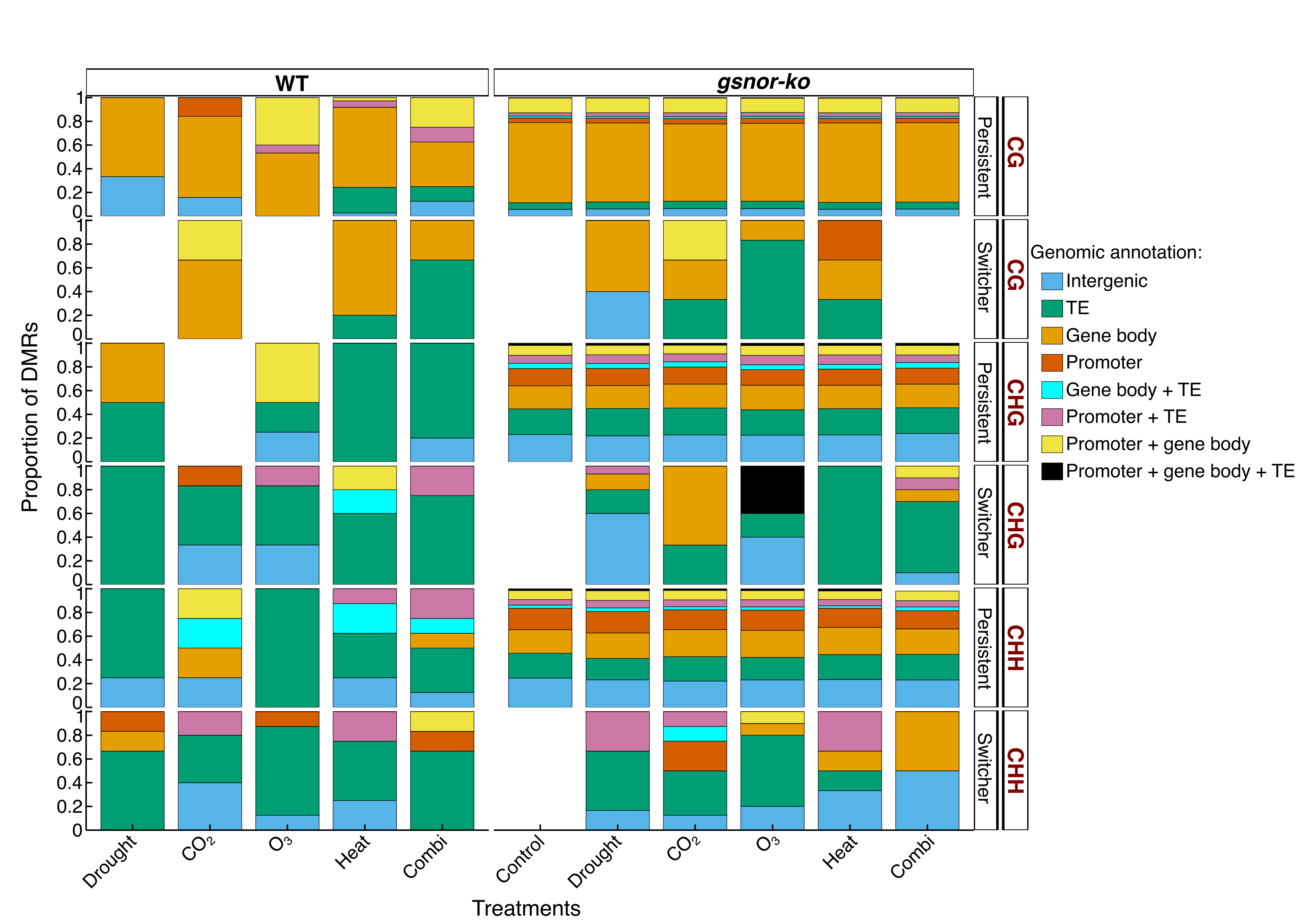

**Supplemental Figure S17 - Genomic annotation of persistent and switcher DMRs across all three sampled generations.** Proportions of persistent and switcher DMRs assigned to genomic annotation classes in WT and *gsnor-ko* plants, separated by treatment lineage and CG, CHG or CHH context. Persistent regions retained the same differential methylation direction at G1, G3 and G5; switchers were detected at all three sampling points but changed direction at least once. All states were defined relative to WT Control within the corresponding generation. Regions were annotated using TAIR10 gene bodies, TEs and promoters, defined as 1 kb upstream of the transcription start site. Colours represent eight mutually exclusive combinations of feature overlaps, including intergenic regions. Each bar shows the distribution of annotations within the indicated DMR class and comparison. Blank positions indicate that no regions were available for that class.

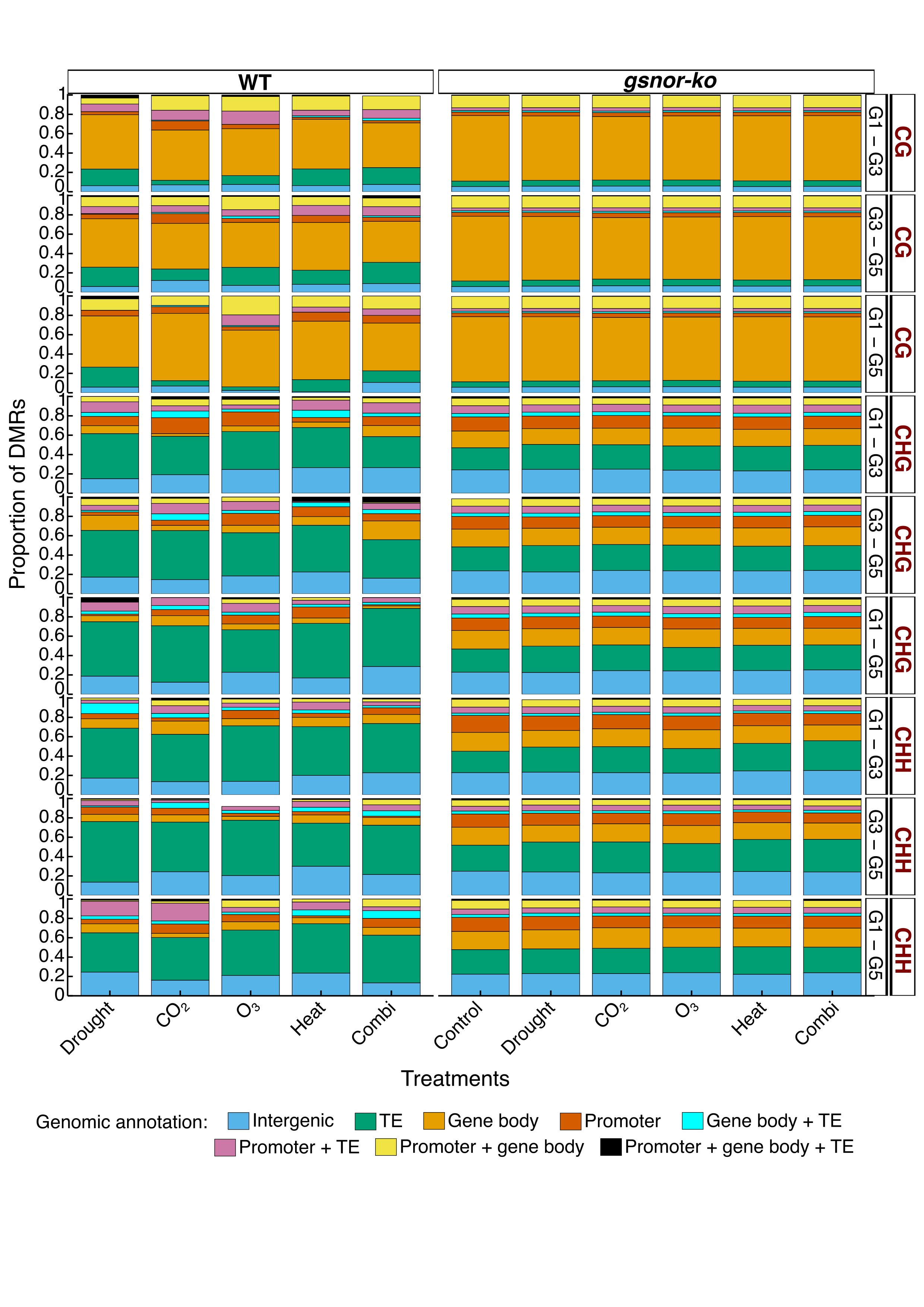

**Supplemental Figure S18 - Genomic annotation of DMRs retaining the same direction between sampled generations.** Proportions of pairwise persistent DMRs assigned to genomic annotation classes for G1–G3, G3–G5 and G1–G5, separated by genotype, treatment lineage and methylation context. Pairwise persistence was defined as differential methylation in the same direction at both indicated sampling points, without requiring persistence in the third sampled generation. Differential methylation states were defined relative to WT Control within each generation. Regions were annotated against TAIR10 gene bodies, TEs and promoters, defined as 1 kb upstream of the transcription start site. Colours indicate eight mutually exclusive feature-overlap classes, including intergenic regions. Proportions were calculated separately within each displayed pairwise persistent DMR set.

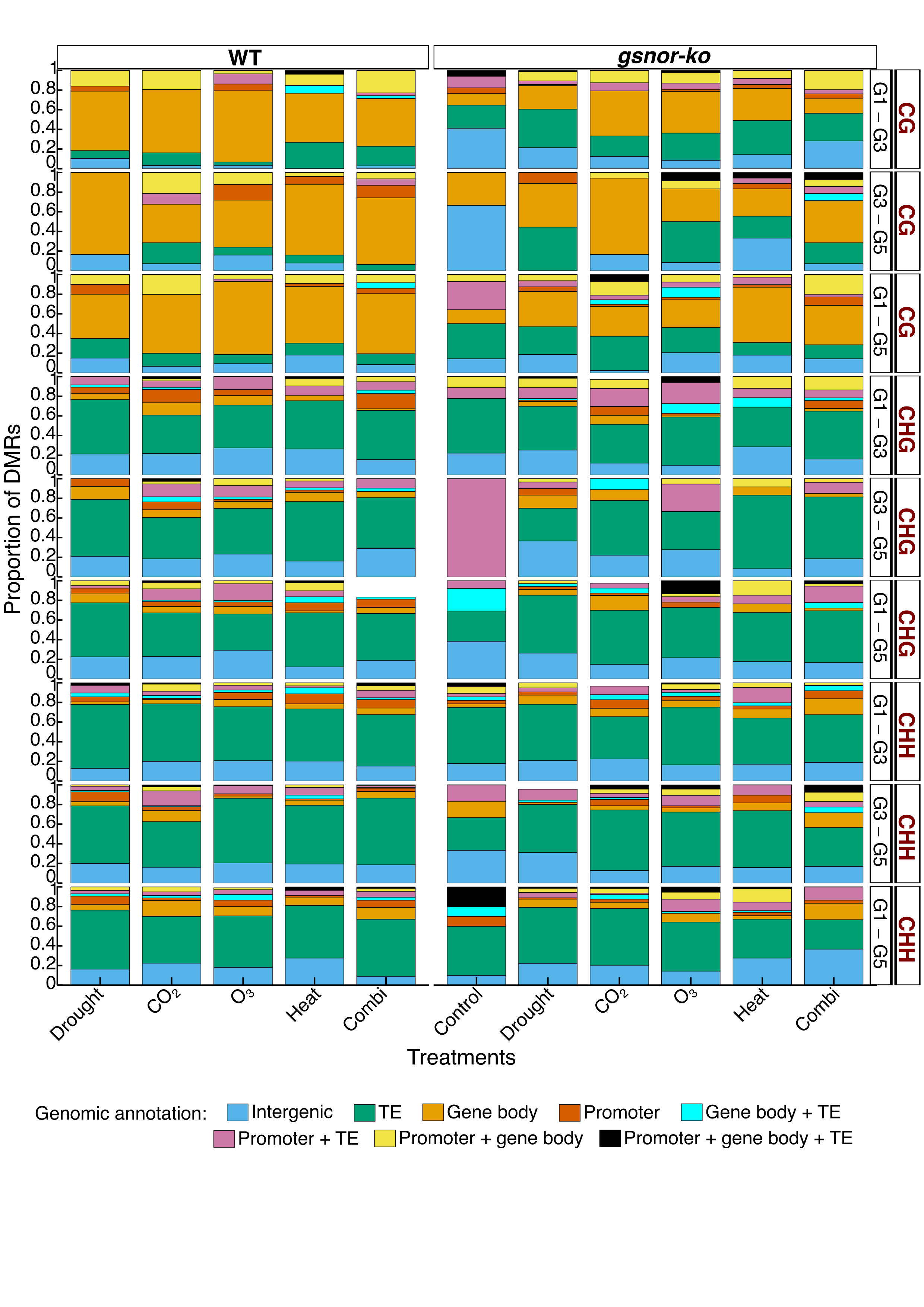

**Supplemental Figure S19 - Genomic annotation of DMRs switching direction between sampled generations.** Proportions of pairwise switcher DMRs assigned to genomic annotation classes for G1–G3, G3–G5 and G1–G5, separated by genotype, treatment lineage and methylation context. Switchers were defined as regions hypermethylated at one indicated sampling point and hypomethylated at the other, relative to the corresponding generation’s WT Control. Classification did not require a particular state in the third sampled generation. Regions were annotated against TAIR10 gene bodies, TEs and promoters, defined as 1 kb upstream of the transcription start site. Colours indicate eight mutually exclusive feature-overlap classes, including intergenic regions. Proportions were calculated separately within each displayed switcher DMR set.

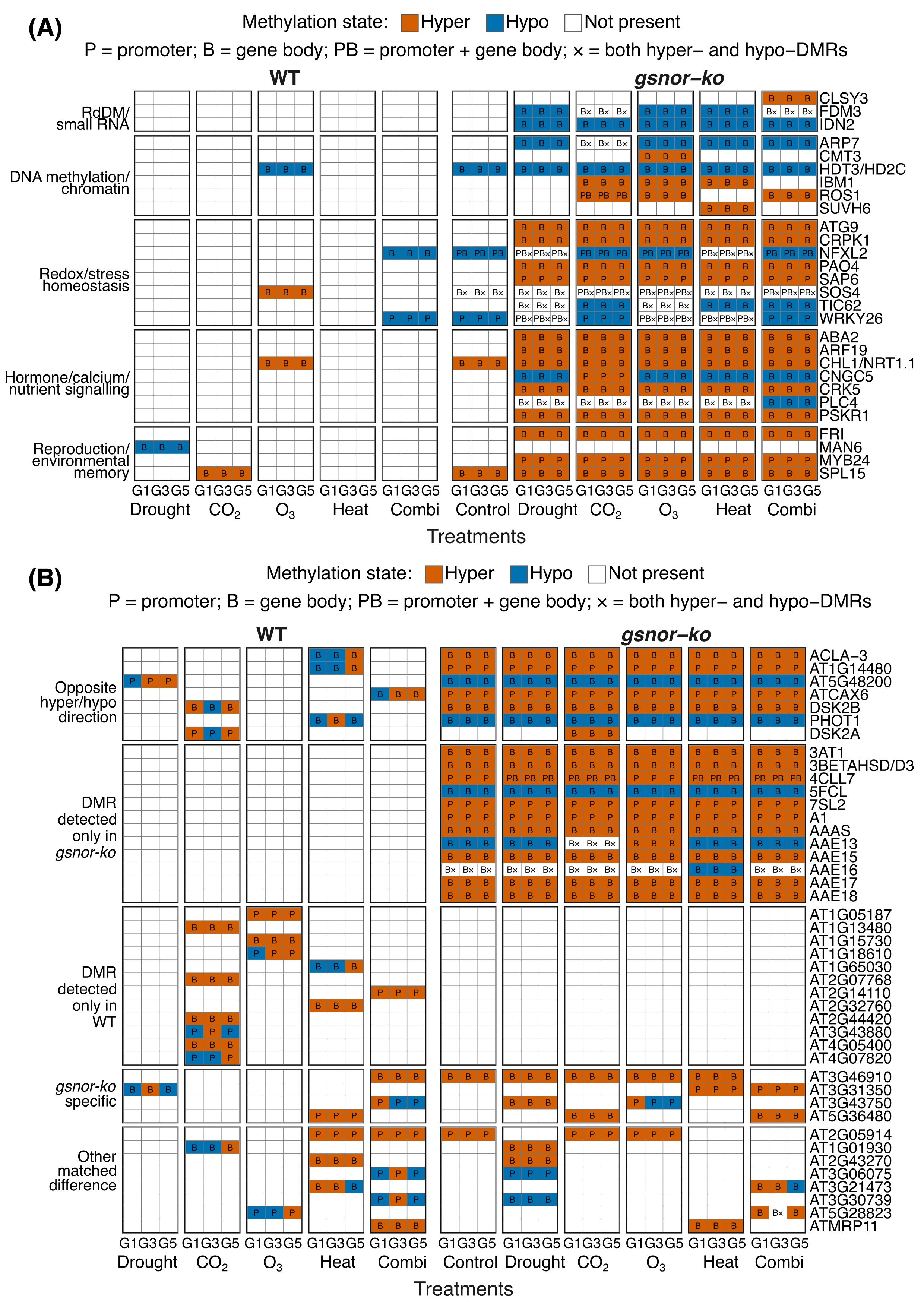

**Supplemental Figure S20 - Expanded candidate-gene methylation patterns and differences between genotypes.** **(A)** Persistent DMR associations for an expanded set of literature-supported candidate genes, grouped by functions related to small-RNA pathways, DNA methylation and chromatin regulation, redox and stress homeostasis, signalling, and reproduction or environmental memory. **(B)** Selected genes grouped according to differences between WT and *gsnor-ko* across matched treatment–generation combinations, including opposite methylation directions, genotype-restricted DMR detection and other differences in methylation state or genomic location. These categories summarise differences across matched cells and do not necessarily indicate absolute genotype specificity across all conditions. Columns within each treatment lineage represent G1, G3 and G5. Orange and blue indicate hypermethylated and hypomethylated DMR associations relative to WT Control within the corresponding generation. White indicates no displayed association, except where × denotes both hypermethylated and hypomethylated DMRs associated with the same gene. P, B and PB indicate promoter, gene-body, and combined promoter/gene-body associations, respectively. Functional grouping and genotype-difference categories are descriptive rather than statistical enrichment results. G5 labels indicate ancestral treatment lineages following two generations under control conditions.

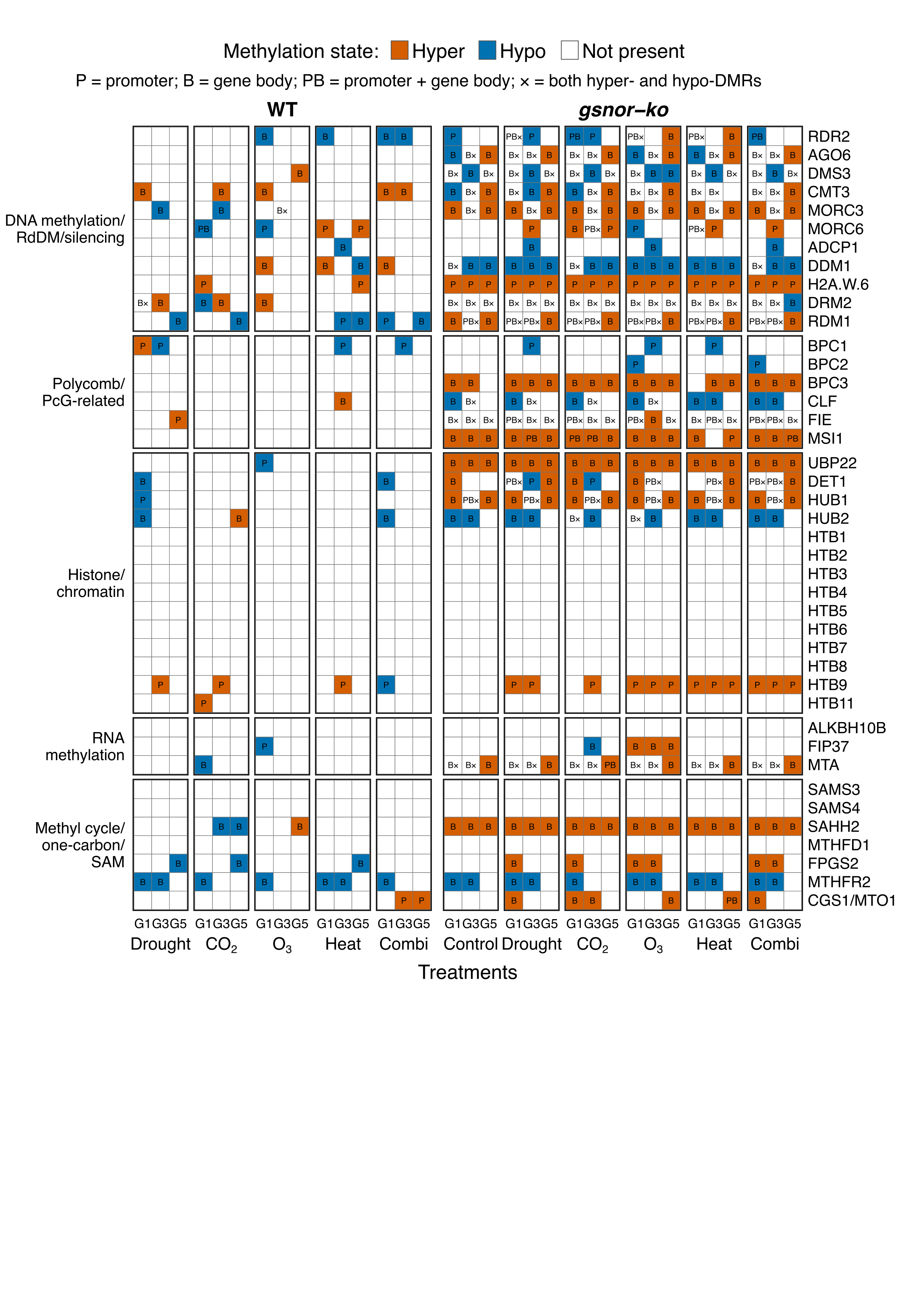

**Supplemental Figure S21 - Generation-specific methylation differences at epigenetic-regulatory genes identified through transcriptome analysis.** Heatmap showing DMR associations for selected genes involved in DNA methylation, RNA-directed DNA methylation (RdDM) and silencing, Polycomb-related regulation, histone and chromatin regulation, RNA methylation, and methyl-cycle or one-carbon metabolism. Candidates were drawn from the associated RNA-seq study. Columns within each treatment lineage represent G1, G3 and G5 in WT and *gsnor-ko* plants. Orange and blue indicate hypermethylated and hypomethylated DMR associations relative to WT Control within the corresponding generation. White indicates no detected association, except where × denotes both hypermethylated and hypomethylated DMRs associated with the same gene. P, B, and PB indicate promoter, gene body, and combined promoter/gene body associations, respectively; promoters were defined as 1 kb upstream of the transcription start site. All generation-specific DMRs were considered, without requiring persistence or switching across all three sampling points. G5 labels indicate ancestral treatment lineages following two generations under control conditions.

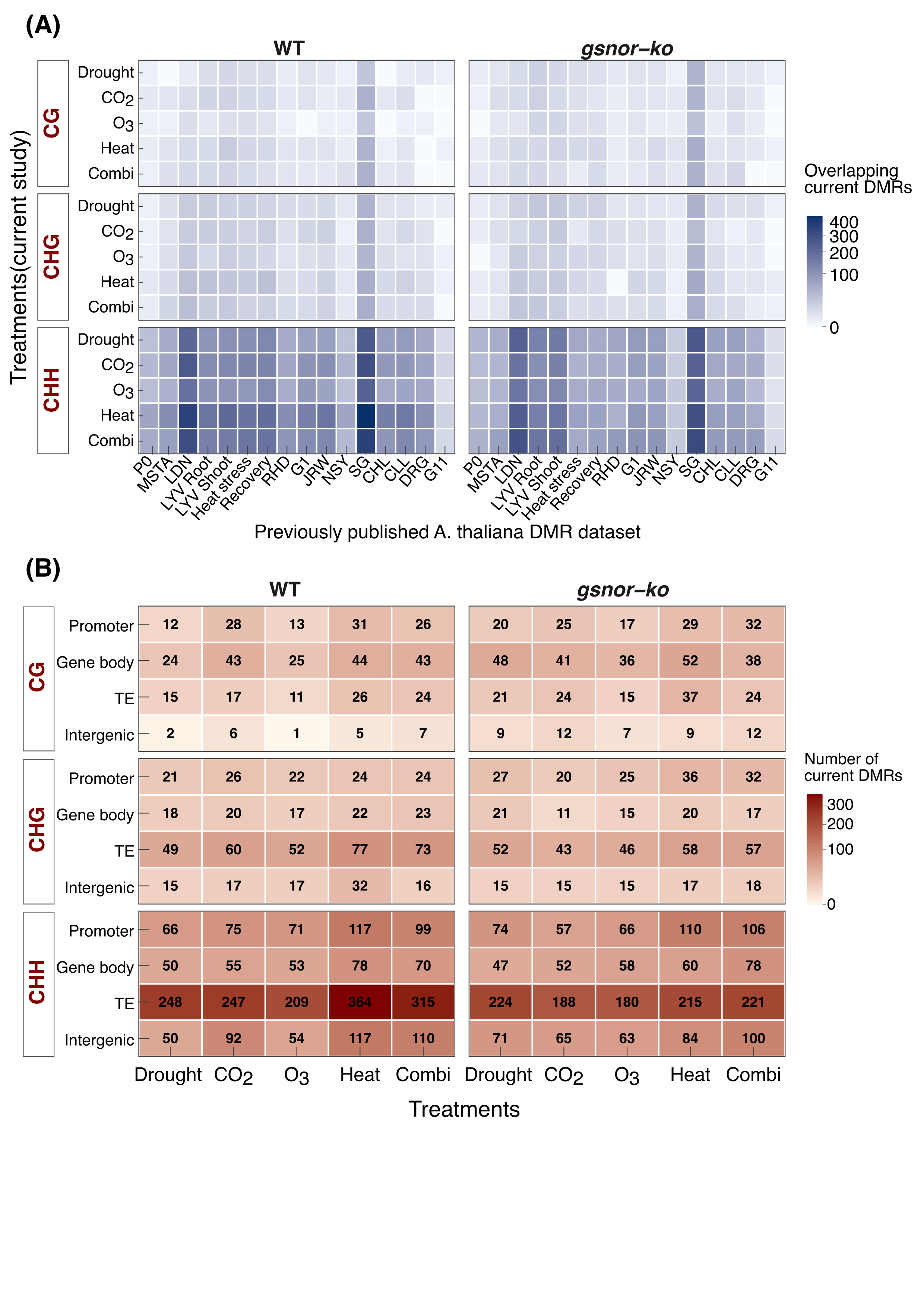

**Supplemental Figure S22 - Overlap of persistent and recurrent DMRs with previously published *Arabidopsis thaliana* stress-methylome datasets.** **(A)** Heatmaps showing the numbers of unique current DMRs overlapping previously reported DMRs, separated by genotype, treatment lineage and methylation context. The persistent/recurrent group includes regions detected in the same direction across G1–G3–G5, regions maintained from G3 to G5, and regions recurrent at G1 and G5. Overlaps required intersecting genomic coordinates within the same methylation context; agreement in methylation direction between studies was not required. Blue shading indicates overlap counts for each published dataset. **(B)** Genomic annotation of current persistent/recurrent DMRs overlapping at least one previous dataset. Numbers and red shading indicate unique current DMR counts within each feature category. Promoter, gene-body and TE annotations are nonexclusive, so a region may contribute to more than one category; intergenic regions overlap none of these features. Current DMRs were defined relative to WT Control within the corresponding generation. Published-dataset identifiers are shown on the horizontal axis in panel A.

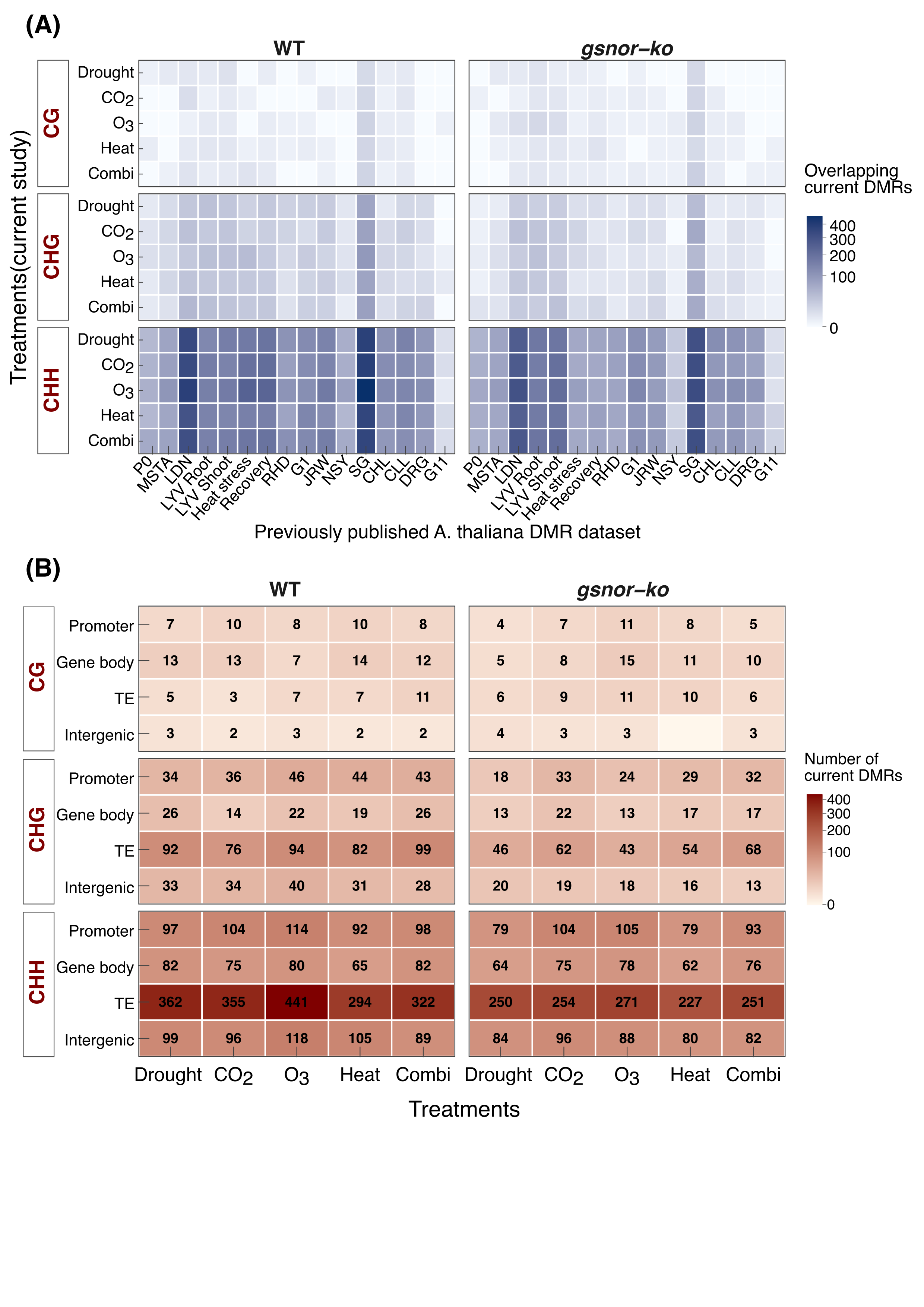

**Supplemental Figure S23 - Overlap of switching DMRs with previously published *Arabidopsis thaliana* stress-methylome datasets.** **(A)** Heatmaps showing the numbers of unique current DMRs classified as switching trajectories that overlap with previously reported DMRs, separated by genotype, treatment lineage, and CG, CHG, or CHH context. Overlaps required intersecting genomic coordinates within the same methylation context; agreement in methylation direction between studies was not required. Blue shading indicates overlap counts for each published dataset. **(B)** Genomic annotation of current switching DMRs overlapping at least one previous dataset. Numbers and red shading indicate unique current DMR counts within promoter, gene-body, TE or intergenic categories. Feature annotations are nonexclusive, except that intergenic regions overlap none of the other three features. Current methylation states were defined relative to WT Control within the corresponding generation. Published-dataset identifiers are shown on the horizontal axis in panel A.

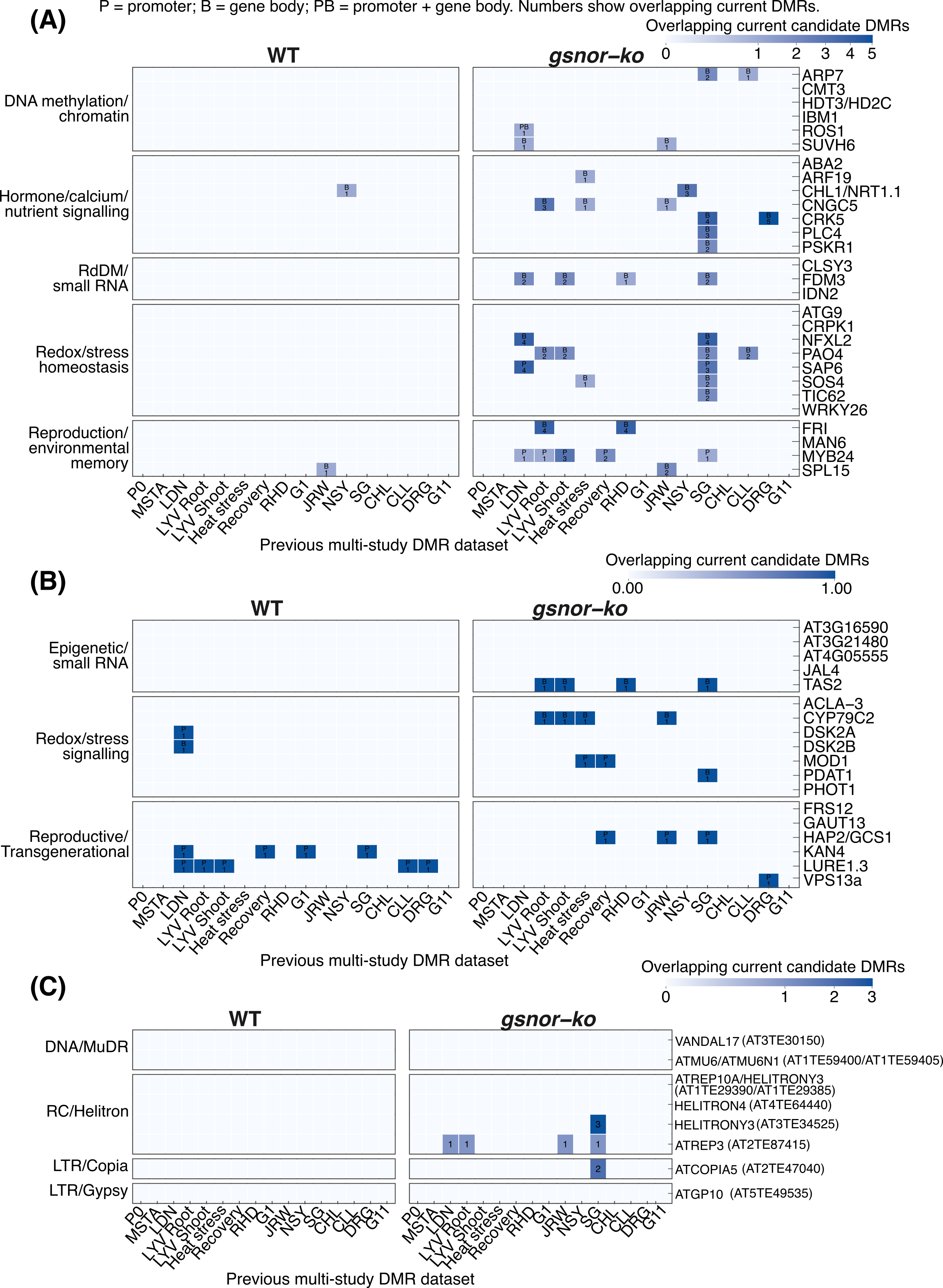

**Supplemental Figure S24 - Independent overlap support for selected gene- and TE-associated DMR candidates.** Heatmaps showing overlaps with previously published *Arabidopsis thaliana* stress-associated DMRs for **(A)** persistent candidate genes, **(B)** switcher candidate genes and **(C)** selected persistent TE candidates. Within each panel, WT is shown on the left and *gsnor-ko* on the right. Rows identify candidates, grouped by functional category or TE superfamily, and columns identify previous datasets. Support required a current candidate-associated DMR to overlap a previous DMR in the same methylation context. Colour intensity and numbers indicate the number of unique overlapping current candidate-associated regions, summarised across the relevant treatment and generation records. P, B and PB indicate promoter, gene-body, and combined promoter/gene-body associations, respectively. Blank cells indicate no detected overlap. Counts represent coordinate-overlap support and do not establish that methylation changes have the same direction or biological effect across studies. Colour scales differ among panels.

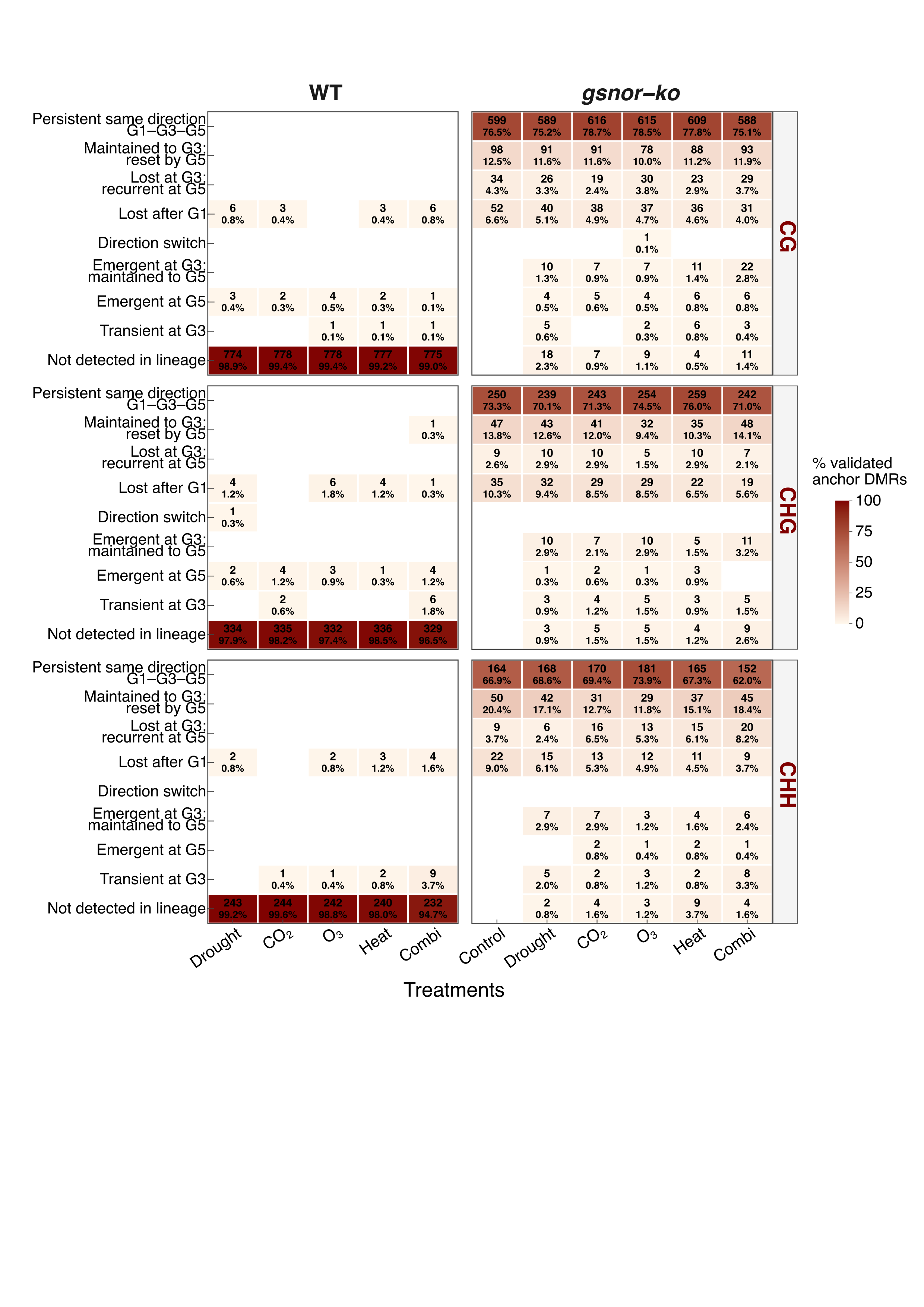

**Supplemental Figure S25 - Generational trajectories of independently supported GSNOR-associated DMRs in WT and mutant lineages.** Heatmaps showing the fate of validated anchor regions across G1, G3 and G5 in WT treatment lineages and *gsnor-ko* control and treatment lineages. Anchors were defined as current G1 *gsnor-ko* Control DMRs overlapping at least one DMR reported by Rudolf *et al.* (2021) in the same hypermethylated or hypomethylated direction. Regions were subsequently tracked across the current lineages and classified according to their detection and direction at the three sampling points. Cell labels show region counts and percentages; shading represents the percentage of the context-specific anchor set. The denominators are 783 CG, 341 CHG, and 245 CHH anchor regions. Persistent regions retained the same direction of differential methylation across all three sampling points. Other categories distinguish loss, recurrence, emergence, transient detection and direction changes. “Not detected in lineage” means that no DMR was detected at any of the three sampling points relative to the corresponding WT Control. Published DMRs lacked individual sequence-context assignments, so validation established coordinate and directional agreement rather than context agreement. G5 treatment labels indicate ancestral exposure followed by two generations under control conditions.

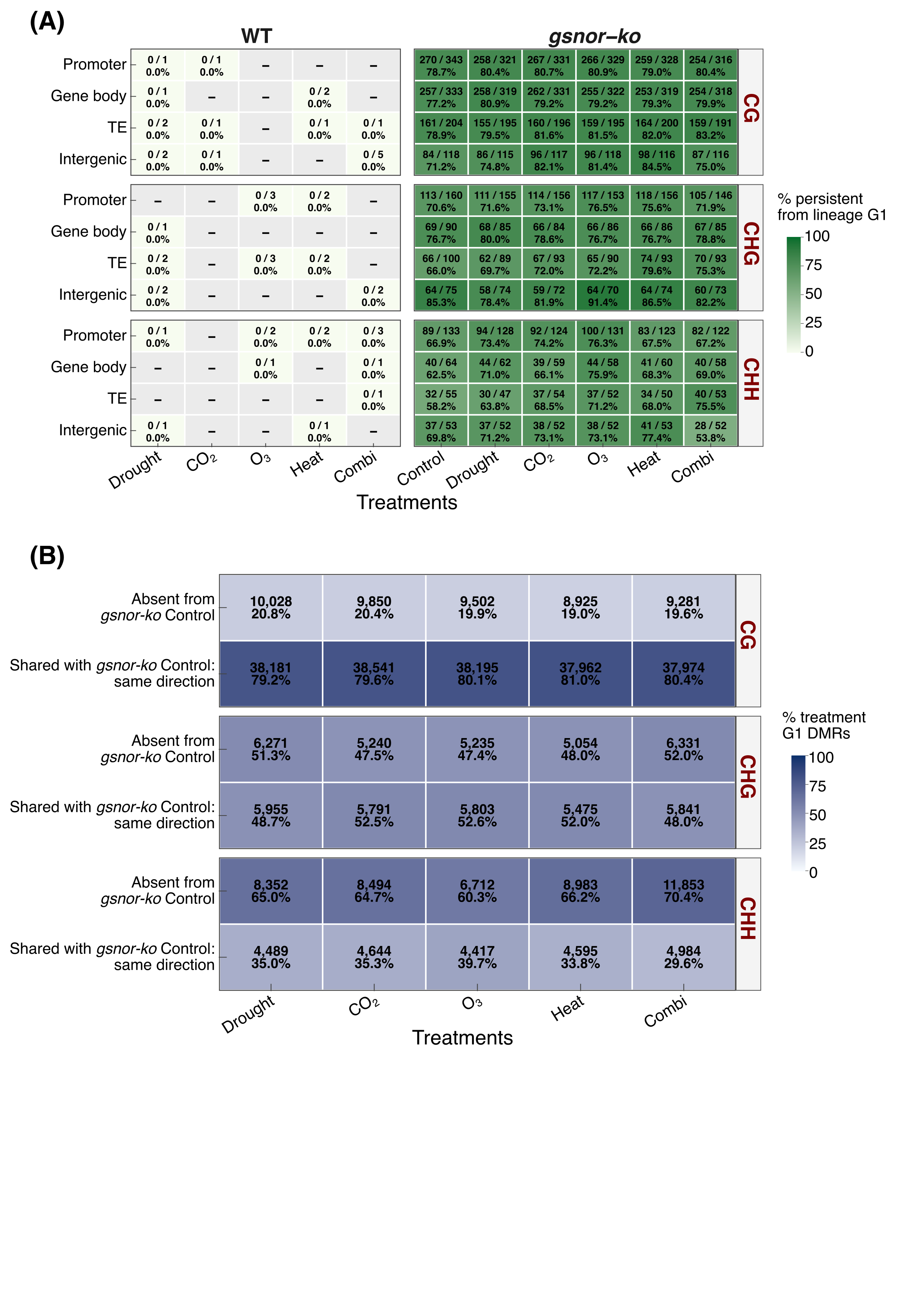

**Supplemental Figure S26 - Annotation-specific persistence of validated regions and separation of mutant baseline from treatment-associated DMRs.** **(A)** Persistence of independently supported anchor regions within promoter, gene-body, TE and intergenic categories, separated by genotype, treatment lineage and methylation context. Anchors were established by coordinate overlap and directional agreement between current G1 *gsnor-ko* Control DMRs and the published DMRs of Rudolf *et al.* (2021). Labels show the number retaining the same direction across G1, G3 and G5 divided by the number detected at that lineage’s G1, followed by the percentage. Green shading represents this percentage. A dash indicates that no anchor regions were detected at lineage G1 in that category. Feature annotations are nonexclusive, except for intergenic regions. **(B)** Classification of the full G1 *gsnor-ko* treatment DMR sets according to their relationship with G1 *gsnor-ko* Control DMRs. Regions were classified as absent from mutant Control or shared with mutant Control in the same methylation direction, using matching sequence context and consensus coordinates. Labels show counts and percentages of the corresponding treatment’s G1 DMR set; blue shading represents these percentages. All original differential methylation states were defined relative to WT Control. Panel B uses the full treatment DMR sets rather than only the independently validated anchors.
